# Metabolic flux and flux balance analyses indicate the relevance of metabolic thermogenesis and aerobic glycolysis in cancer cells

**DOI:** 10.1101/2021.11.16.468557

**Authors:** Nobuyuki Okahashi, Tomoki Shima, Yuya Kondo, Chie Araki, Shuma Tsuji, Akane Sawai, Hikaru Uehara, Susumu Kohno, Hiroshi Shimizu, Chiaki Takahashi, Fumio Matsuda

## Abstract

Adenosine triphosphate (ATP) regeneration by substrate-level phosphorylation is a general feature of cancer metabolism, even under normoxic conditions (aerobic glycolysis). However, it is unclear why cancer cells prefer inefficient aerobic glycolysis over the highly efficient process of oxidative phosphorylation for ATP regeneration. To investigate the metabolic principles underlying aerobic glycolysis, we performed ^13^C-metabolic flux analysis of 12 cultured cancer cell lines and explored the metabolic constraints required to reproduce the results using *in silico* metabolic simulations. We found that the measured flux distribution can be reproduced by maximizing the ATP consumption in the flux balance analysis considering a limitation of metabolic heat dissipation (enthalpy change). It suggests that aerobic glycolysis may be preferable because metabolic heat production during one mol of ATP regeneration by aerobic glycolysis was less than that produced by oxidative phosphorylation (OXPHOS). Consistent with the simulation, OXPHOS inhibition induced metabolic redirection to aerobic glycolysis while maintaining the intracellular temperature. Furthermore, the dependency on aerobic glycolysis was partly alleviated upon culturing at low temperatures. Our data suggest that metabolic thermogenesis is an important factor in understanding aerobic glycolysis in cancer cells and that an advantage of aerobic glycolysis is the reduction in metabolic heat generation during ATP regeneration.

## Introduction

Adenosine triphosphate (ATP) regeneration is one of the most important metabolic processes and is required to maintain various cellular functions. Generally, normal human cells efficiently regenerate ATP by oxidative phosphorylation (OXPHOS) through the electron transport chain (ETC) (32 ATP/glucose, denoted as the glucose → TCA cycle). In contrast, cancer cells depend on substrate-level phosphorylation for ATP regeneration, even under conditions of oxygen availability (2 ATP/glucose, aerobic glycolysis) ^1^. This altered metabolic state is ascribed to the reduced ETC activity that triggers the metabolic rewiring toward aerobic glycolysis to meet the large ATP demand for active cell proliferation in cancer cells ^2^. However, ETC deficiency depends on the type of cancer cell line as certain cell types activate ETC and more than 50% of ATP regeneration is attributed to OXPHOS ^3^. Moreover, cancer cells commonly catabolize glutamine as a carbon source to regenerate ATP by OXPHOS (glutaminolysis) ^4, 5^. Thus, the rationale for employing aerobic glycolysis for ATP regeneration is far from being understood, although several possible roles have been proposed ^6^.

## Results

### Total ATP regeneration flux is not correlated with cell proliferation rates

In this study, we performed ^13^C-metabolic flux analysis (^13^C-MFA) of 12 cancer cell lines and *in silico* metabolic simulations. These cell lines were selected based on the diversity of their expression patterns for genes associated with central carbon metabolism to obtain a global view of energy metabolism ^7^ (**Fig. S1**). Their ability to grow in Dulbecco’s modified Eagle’s medium (DMEM) was considered to reduce the bias derived from the culture conditions. The specific cell proliferation rate, cell adhesion area, and diameter of trypsinized cells were measured as visible phenotypes of the cell cultures (**Table S1**).

Metabolic flux distributions allowed quantitative comparison of flux levels among pathways ^8, 9^ (**Figs. 1A and S2, Table S2, Data S1**). Details of the procedure for ^13^C-MFA are provided in the supplementary Text (**Text S1**). ATP was mainly regenerated by three metabolic pathways in the 12 cancer cell lines: aerobic glycolysis (red arrow in **Fig. 1A**), glutaminolysis (blue arrow), and glucose → TCA cycle (gray arrow). In the case of MCF-7 cells, the metabolic flux level in glycolysis, represented by the glyceraldehyde-3-phosphate dehydrogenase (GAPDH) reaction, was 1104 nmol (10^6^ cells)^-1^ h^-1^. The flux levels of glutamate dehydrogenase (GLUDH, representing the glutaminolysis flux) and the isocitrate dehydrogenase (IDH) reaction (representing the glucose → TCA cycle flux) were less than 10% of the glycolytic flux, that were 19 and 25 nmol (10^6^ cells)^-1^ h^-1^, respectively (**Fig. 1A**). All 12 cell lines demonstrated similar trends, that is, the flux through aerobic glycolysis was generally higher than that through glutaminolysis and the glucose → TCA cycle (**Fig. S3**). However, significant variations were observed in the fluxes of GAPDH, GLUDH, and IDH reactions among the cell lines (**Fig. S3**). Among these, the glucose → TCA cycle flux was not universal because the metabolic flux levels of the IDH reaction were close to zero in HepG2 and MIA PaCa-2 cells (**Fig. S3**). Principal component analysis revealed no definite correlation between the metabolic flux distributions and cell origins (**Fig. 1B**), suggesting that the variations may be due to other biological contexts in each cell line.

**Figure 1.**
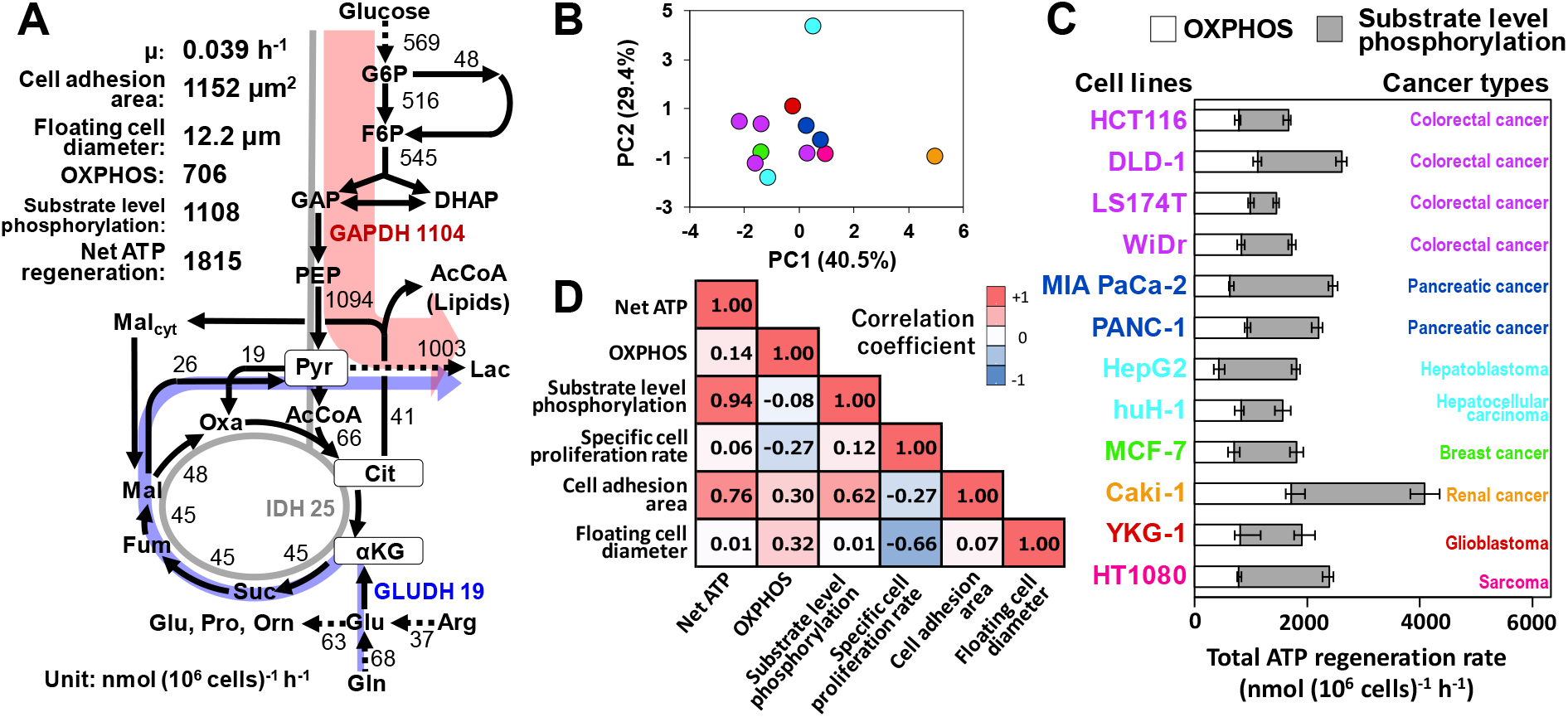
Variation in the metabolic flux distribution among 12 cancer cell lines and their correlation with the phenotypes. **(A)** Metabolic flux distribution of MCF-7 as determined by ^13^C-MFA. **(B)** The score plot of the principal component analysis using the metabolic flux distributions of 12 cell lines. The color legend is identical to that of the next panel. **(C)** Variations in ATP regeneration rates by substrate-level phosphorylation and OXPHOS. Error bars represent 95% confidence intervals. **(D)** Correlation between ATP regeneration rates and visible cell phenotypes. Spearman’s rank order correlation coefficients are depicted in the heat map (*n*=12).

Net ATP regeneration rates within the central carbon metabolic network were calculated from the metabolic flux distributions (**Fig. 1C**). For instance, the net ATP regeneration rate in MCF-7 cells was 1815 nmol (10^6^ cells)^-1^ h^-1^, which varied 2.8 times among the 12 cell lines. An approximate estimate indicated that the net ATP regeneration rates were higher than those of most non-cancerous cells (See Supplementary **Text S2** for details). Moreover, the ATP regeneration rate in MCF-7 cells showed that 39% of ATP was regenerated by OXPHOS through the ETC (**Fig. 1C**). OXPHOS commonly occurred in all 12 cancer cell lines, suggesting that this mechanism is still mandatory for ATP regeneration in cancer cells. Consistent with this observation, the cell lines maintained their mitochondrial membrane potential (**Fig. S4**). However, a large variation was observed in the OXPHOS-dependent ATP regeneration rate, ranging from 24% (HepG2) to 68% (LS174T) of the net ATP regeneration rate (**Fig. 1C**) ^10^. This variation is likely to be derived from the distinct ETC activities of each cell line depending on the biological context. Thus, it is reasonable to assume that the ETC must work within an optimal capacity as a loss of mitochondrial membrane potential by low ETC activity triggers cell apoptosis ^11, 12^ and an ETC overload excessively generates reactive oxygen species, which also induces cell apoptosis ^13^.

Given that cells maintain a rapid turnover of ATP, it can be assumed that an equal amount of regenerated ATP must be consumed for some cellular functions, such as proliferation. However, no correlation was observed between ATP regeneration rates and specific cell proliferation rates (*r* = 0.06, **Figs. 1D and S5**), suggesting the presence of other major ATP demands in cultured cancer cells. The correlation analysis indicated that the cell adhesion area on the culture plates was positively correlated with the net ATP regeneration rates (*r* = 0.76, **Figs. 1D and S5**). This trend was also observed when an outlier (Caki-1) was removed from the dataset (*r* = 0.68; **Fig. S6**). Moreover, no correlation was observed between the cell adhesion area and diameter of the trypsinized floating cells (*r* = 0.07, **Fig. 1D**). Thus, the cell adhesion area is a phenotype of cancer cells that is distinct from the cell volume. Notably, the adhesion between the cell and the extracellular matrix mediated by cadherin promotes cancer cell invasion in conjunction with metabolic changes to produce the required amount of ATP ^14^. Actin and myosin utilize ATP to regulate tumor cell adhesion and migration ^15, 16^. Taken together, these results indicate that cancer cell metabolism is not shaped by a proliferation-oriented mechanism.

### *In silico* simulation indicates that aerobic glycolysis is required to reduce metabolic heat

To elucidate the principle underlying the coordination of cancer metabolism, we explored the metabolic constraints required to reproduce the measured flux distributions using *in silico* metabolic simulations. We performed flux balance analysis (FBA) using a genome-scale model of human metabolism (RECON2) ^17^ (**Data S2**). FBA requires an objective function to simulate the intracellular metabolic flux distribution using a linear programming method ^18^. Although maximal biomass production is commonly used as an objective function to simulate bacterial metabolism, a suitable objective function for simulating cancer cell metabolism remains under investigation ^19^. In this study, the ^13^C-MFA results suggested that the cultured cancer cells consumed a large amount of ATP to maintain certain cellular functions other than proliferation (**Fig. 1D**). Thus, the maximal ATP consumption was employed as the objective function in the *in silico* analysis.

The FBA also requires constraints on the solution space (upper and lower boundaries of the metabolic flux vector) ^20^. Here, we performed FBA to determine additional constraints that drive cell metabolism toward aerobic glycolysis and glutaminolysis for ATP regeneration, as revealed in the ^13^C-MFA (**Fig. 1**). The flux distribution of MCF-7 cells was chosen as an example for reproduction by FBA (**Fig. 2A**). The relatively minor metabolic fluxes, such as those related to biomass synthesis as well as the uptake and production of amino acids other than glutamine, were constrained at the measured values of MCF-7 (all constraints for the influx and efflux are presented in **Table S3**). In contrast, no specific constraints were imposed on the uptake rates of glucose, glutamine, and oxygen and the excretion rates of lactate, carbon dioxide, and water.

**Figure 2.**
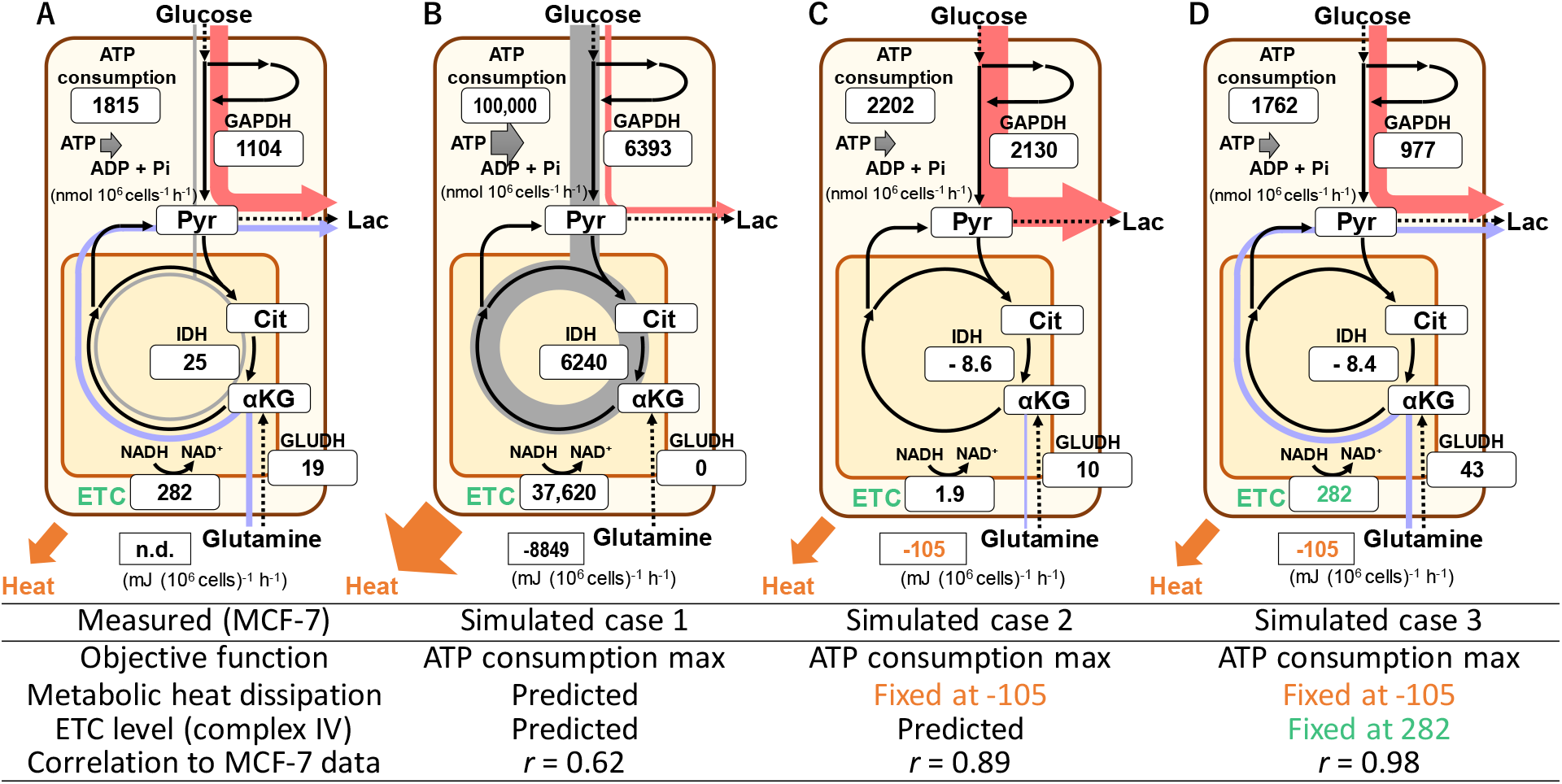
Metabolic heat dissipation and ETC activity govern cancer cell metabolism. *In silico* simulation of metabolism was performed using the FBA with a genome-scale model of human metabolism (RECON2). Metabolic flux levels of the key reactions of aerobic glycolysis (red), glucose → TCA cycle (gray), and glutaminolysis (blue) are displayed in the boxes. The maximum ATP consumption was employed as an objective function. Additional constraints are represented by yellow and green numbers. **(A)** Measured metabolic flux distribution in MCF-7 cells. **(B-D)** Results of FBA **(B)** without additional constraints, **(C)** with fixed metabolic heat dissipation, and **(D)** with fixed metabolic flux dissipation and ETC flux.

The FBA results without additional constraints showed that considerable amounts of glucose and glutamine were consumed until the ATP regeneration rate reached the upper limit (**Fig. 2B, Data S3**). The predicted metabolic flux distribution mainly exhibited OXPHOS-dependent metabolism through the glucose → TCA cycle and a low correlation coefficient with that observed in MCF-7 cells (*r* = 0.62). This indicated that additional constraints are required to reproduce the measured value for MCF-7 cells.

In this study, we considered various thermodynamic parameters that govern cell metabolism ^21^. For example, the entire stoichiometry of aerobic glycolysis (glucose → 2 lactate) indicates that the enthalpy change (Δ_r_*H*’°), or metabolic heat generation, per 1 mole of ATP regeneration by aerobic glycolysis is −55.0 kJ (mol ATP)^-1^ (**Table S4**) ^22^. Similarly, the enthalpy change in the glucose → TCA cycle reaction was −88.2 kJ (mol ATP)^-1^ (**Table S4**). Thus, it is expected that if the enthalpy change is limited, aerobic glycolysis would be preferentially employed for ATP regeneration in cells. Thermogenesis has long been observed in patients with breast cancer ^23^. The preference for aerobic glycolysis was not observed for other parameters, such as entropy and Gibbs free energy change (**Table S4**). Thus, the genome-scale metabolic model was modified to determine the levels of cellular dissipation of enthalpy change from the FBA results (**Table S5 and Data S3**) ^22^.

The FBA was performed by fixing the Δ_r_*H*’° level at an arbitrary value as an additional constraint. We found that when the Δ_r_*H*’° level was set at −105 mJ (10^6^ cells)^-1^ h^-1^, the predicted metabolic flux distribution was more similar to the measured MCF-7 data (**Fig. 2C**, *r* = 0.89) because aerobic glycolysis was activated for ATP regeneration, as expected.

In addition to metabolic heat, the ETC activity was applied as an additional constraint because the level of ETC flux in the above simulation was considerably low (**Fig. 2C**), which was inconsistent with the measured data (**Figs. 2A and S4**). Thus, FBA was performed by fixing the metabolic flux level of complex IV of the ETC at the measured value of MCF-7 cells (282 nmol (10^6^ cells)^-1^ h^-1^). The predicted metabolic flux distribution successfully reproduced ATP regeneration by both aerobic glycolysis and glutaminolysis, and its correlation with the measured flux distributions was *r* = 0.98 (**Fig. 2D, Fig. S7, Data S3**). However, ATP regeneration by the glucose → TCA cycle did not occur in the predicted distribution (**Fig. 2D**), which was inconsistent with the ^13^C-MFA result (**Fig. 2A**). This is because the metabolic heat dissipation per 1 mole of ATP regeneration by glutaminolysis is lower than that of the glucose → TCA cycle reaction, although the detailed −Δ_r_*H*’° level cannot be calculated by simple material balance equations. Notably, the glucose → TCA cycle pathway is not universal, as mentioned above (**Fig. S3**). The FBA showed that similar ATP regeneration occurred by both aerobic glycolysis and glutaminolysis in all 12 cell lines (**Fig. S7 and Data S3**). The simulation of cancer cell metabolism suggested that the ETC activity and metabolic heat generation are possible constraints of cancer cell metabolism.

### ETC inhibitor treatment induces metabolic rewiring toward aerobic glycolysis while maintaining the intracellular temperature

The metabolic simulation indicates that the downregulation of ETC activity will be compensated by the elevation of aerobic glycolysis while maintaining the heat dissipation level, which corresponds to a metabolic shift from the level in **Fig. 2D** to that in **Fig. 2C**. To test this hypothesis, the MCF-7 cells were treated with various ETC inhibitors. Rotenone treatment (100 nM) partially inhibited cell proliferation by 25% after 24 h (data not shown). The time-course data showed that cell proliferation was temporarily halted by 6 h and then recovered between 6 and 24 h (**Fig. 3A**). The recovery in growth was accompanied by the activation of aerobic glycolysis as the specific rates for glucose consumption and lactate production between 12 and 24 h increased by 1.2-fold with rotenone treatment (**Fig. 3B**).

**Figure 3.**
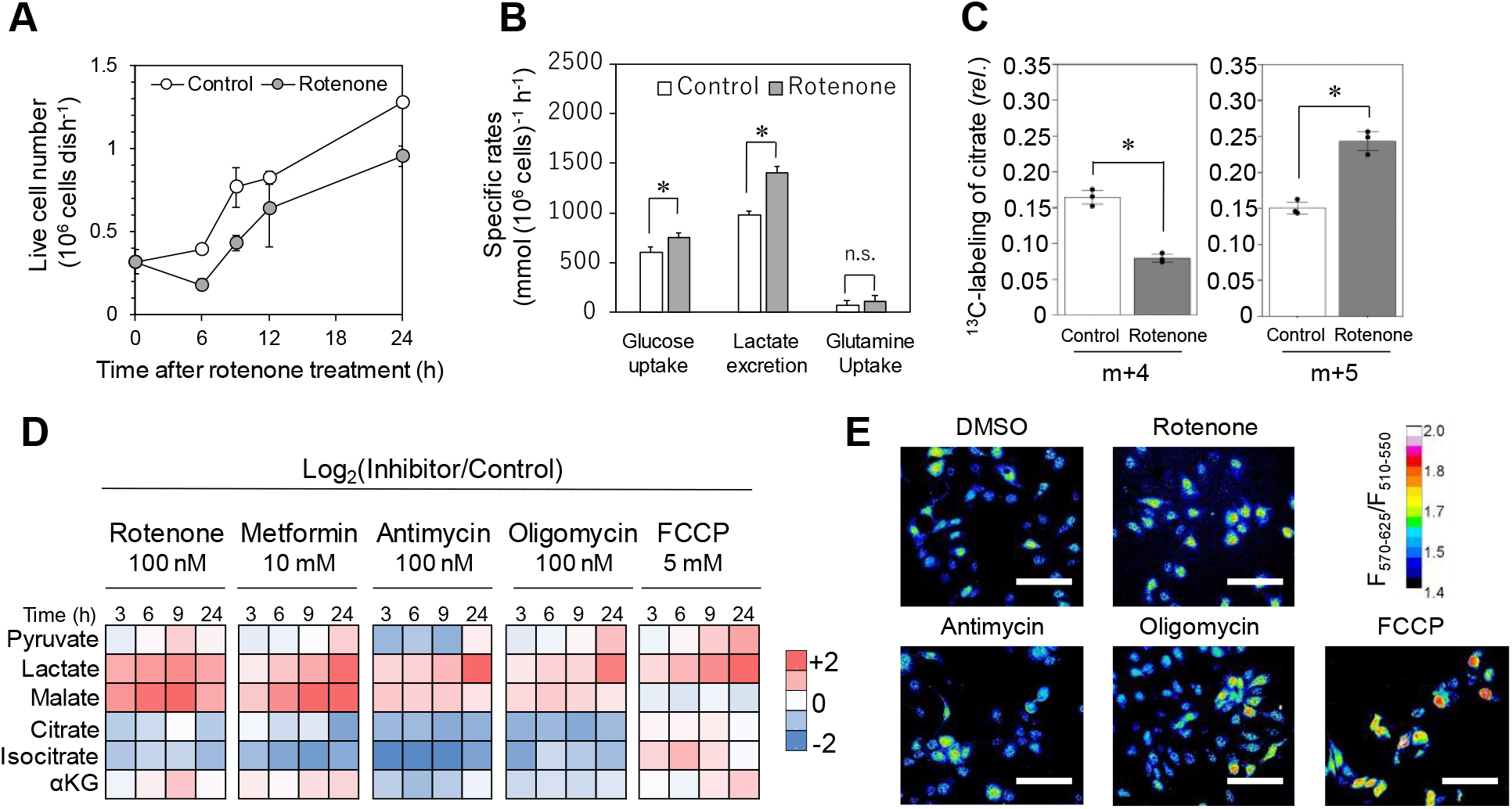
ETC inhibitor treatment induces metabolic rewiring toward aerobic glycolysis while maintaining the intracellular temperature. **(A)** Proliferation of MCF-7 cells after 100 nM rotenone treatment. **(B)** Specific rates for glucose and glutamine uptake and lactate excretion between 12 and 24 h of culturing. Asterisks indicate the statistical significance relative to the control, as determined by a two-sided Student’s *t*-test (*p* < 0.05). **(C)** Mass isotopomer distribution of citrate in MCF-7 cells in [U-^13^C]glutamine tracing. **(D)** Metabolites of MCF-7 cells treated with OXPHOS inhibitors. Relative abundances of metabolites (Log_2_(fold change)) are shown in the heatmap. All data are represented as mean ± SD (*n*=3). **(E)** Intracellular temperature of MCF-7 treated by ETC-inhibitors for 24 h. Intracellular temperatures of MCF-7 cells treated with DMSO, 100 nM rotenone, 100 nM antimycin, 100 nM oligomycin, and 5 mM FCCP for 24 h were measured using a cellular thermoprobe. The color scale represents the fluorescence intensity ratio (570-625/510-550 nm), which corresponds to the intracellular temperature.

To investigate the switching point of the metabolic pathway, ^13^C-tracer and metabolome analysis was performed. Tracer analysis using [U-^13^C]glutamine revealed that the relative abundance of the m+4 isotopomer of citrate, which is produced from acetyl-CoA and oxaloacetate, was reduced after rotenone treatment, whereas that of m+5 citrate produced by reductive glutamine metabolism increased (**Fig. 3C**). The labeling pattern indicated a reduction in the flux of the glucose → TCA cycle in the rotenone-treated MCF-7 cells. The metabolic shift is reasonable because MCF-7 cells used the glucose → TCA pathway (**Fig. 1A**) although the use of this pathway could not be reproduced by metabolic simulation (**Fig. 2D**). Metabolome analysis showed that rotenone treatment induced an accumulation of malate and lactate, while also decreasing the citrate and isocitrate levels (**Fig. 3D**). A similar metabolic profile was commonly induced by other ETC inhibitors, antimycin and oligomycin, but not by the mitochondrial uncoupler, carbonyl cyanide 4-(trifluoromethoxy)phenylhydrazone (FCCP) (**Fig. 3D**). These results suggest that the downregulation of the entry point of the TCA cycle also contributes to a metabolic shift from the glucose → TCA cycle to aerobic glycolysis.

To evaluate the effect of changes in heat generation by metabolic redirection, the intracellular temperature of cells treated with ETC inhibitors was measured using a celluler thermoprobe. The temperature increase caused by the FCCP-induced decoupling was accurately captured (**Fig. 3E**). In contrast, treatment with rotenone and other ETC inhibitors, including antimycin and oligomycin, did not significantly change the intracellular temperature, demonstrating that aerobic glycolysis was elevated while maintaining the heat dissipation level (**Fig. 3E**). The metabolic simulation results indicate that a reduction in ATP regeneration by the restriction of ETC activity was circumvented by the activation of aerobic glycolysis within the heat dissipation capacity, supporting the validity of the metabolic model considering enthalpy change.

### Metabolic rewiring toward aerobic glycolysis is dependent on temperature

The above observations suggest that cancer cells rewire their metabolism to control the heat dissipation level. The glucose → TCA cycle pathway is preferable under lower and higher temperature conditions to maintain a constant intracellular temperature. To test this assumption, cells were cultured at control (37 ℃), low (36 ℃), and high (38 ℃) temperatures. A comparison between the glucose and lactate consumption for 18 h showed that there was a limited effect on the glucose consumption (**Fig. 4A**), whereas lactate production tended to decrease and increase at 36 and 38 ℃, respectively (**Fig. 4B**). A significant increase in lactate production was observed in both cell lines at 38 ℃ (*t*-test, *n*=3, α=0.05). In contrast, a significant decrease in lactate production was observed in five cell lines at 36 ℃. The results indicated that the metabolic shift toward the glucose → TCA cycle pathway was temperature-sensitive.

**Figure 4.**
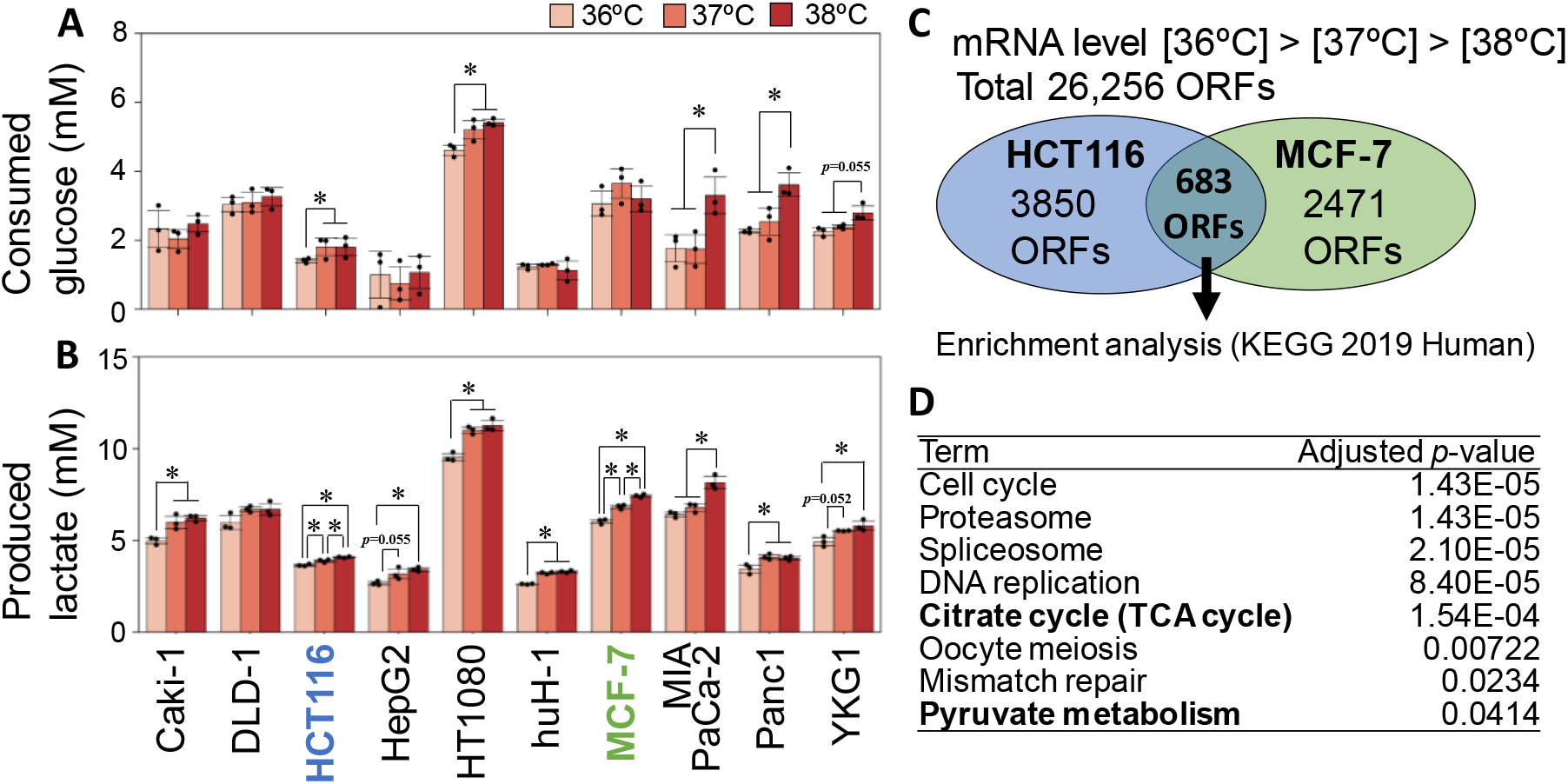
Culture temperature induces metabolic rewiring by altering gene expression. Cancer cells were cultured under control (37 ℃), low (36 ℃), and high (38 ℃) temperatures. Concentrations of **(A)** glucose and **(B)** lactate in the culture medium at 18 h. Data are represented as mean ± SD (*n*=3). Asterisks indicate the statistical significance of concentrations under test conditions as determined by one way analysis of variance (ANOVA) using the Tukey-Kramer method (\**p* < 0.05). **(C)** Venn diagram of temperature-responsive genes. RNA-Seq of MCF-7 and HCT116 cells at 24 h identified 3154 and 4533 genes expressed with fold changes in the following order: [36 ℃]>[37 ℃]>[38 ℃]. **(D)** An over-represented analysis of gene categories of 683 common genes by Enricher using the KEGG 2019 Human dataset ^24^.

Gene expression data were obtained from MCF-7 and HCT116 cells, which indicated a temperature-dependent increase in lactate production, as representative cell lines (*n*=1, **Table S6, Data S4**). We selected a total of 3154 and 4533 open reading frames (ORFs), whose expression levels were altered in MCF-7 and HCT116 cells in the following order: [36 ℃]>[37 ℃]>[38 ℃]. A total of 683 ORFs were common in the MCF-7 and HCT116 datasets (**Fig. 4C, Data S5**), which was significantly larger than the expected number of overlapping ORFs (545 ORFs, χ^2^ test, *p*-value =2.0 × 10^-9^). An over-representation analysis was performed for the 683 ORFs using the Kyoto Encyclopedia of Genes and Genomes (KEGG) 2019 human dataset ^24^. The results revealed that ORFs in the “TCA cycle” category (*PDHA1, MDH2, IDH3G, IDH2, IDH3B, OGDHL, DLAT,* and *SDHD*) were enriched in 683 ORFs (**Fig. 4D, Table S7**). Notably, *PDHA1* and *DLAT* encode subunits of the pyruvate dehydrogenase complex, which is located at the entry point of the TCA cycle; this was also indicated in the aforementioned metabolome analysis (**Fig. 3D**). No functional category was found in the ORF list that exhibited a reversed expression pattern in the following order: [36 ℃]<[37 ℃]<[38 ℃] (data not shown). These results suggest that metabolic rewiring from the glucose → TCA cycle to aerobic glycolysis is dependent on temperature-induced changes in the metabolic thermogenesis.

## Discussion

This study reaffirms that thermogenesis is an important aspect of cellular metabolism, particularly when cancer cells require large amounts of ATP. The effects of thermogenesis on cancer cell metabolism are shown in **Figure 5**. In non-cancerous cells, ATP is supplied by the glucose → TCA cycle pathway through OXPHOS, which efficiently regenerates ATP (**Fig. 5A**). Metabolic heat dissipation is maintained within the heat homeostasis. The ATP demand is elevated during cancer development as ATP is required for cancer-specific features, such as rapid cell proliferation, tumor cell adhesion, and migration. The ^13^C-MFA results suggested that a larger amount of ATP was utilized for cell adhesion rather than cell proliferation (**Figs. 1C and D**).

**Figure 5.**
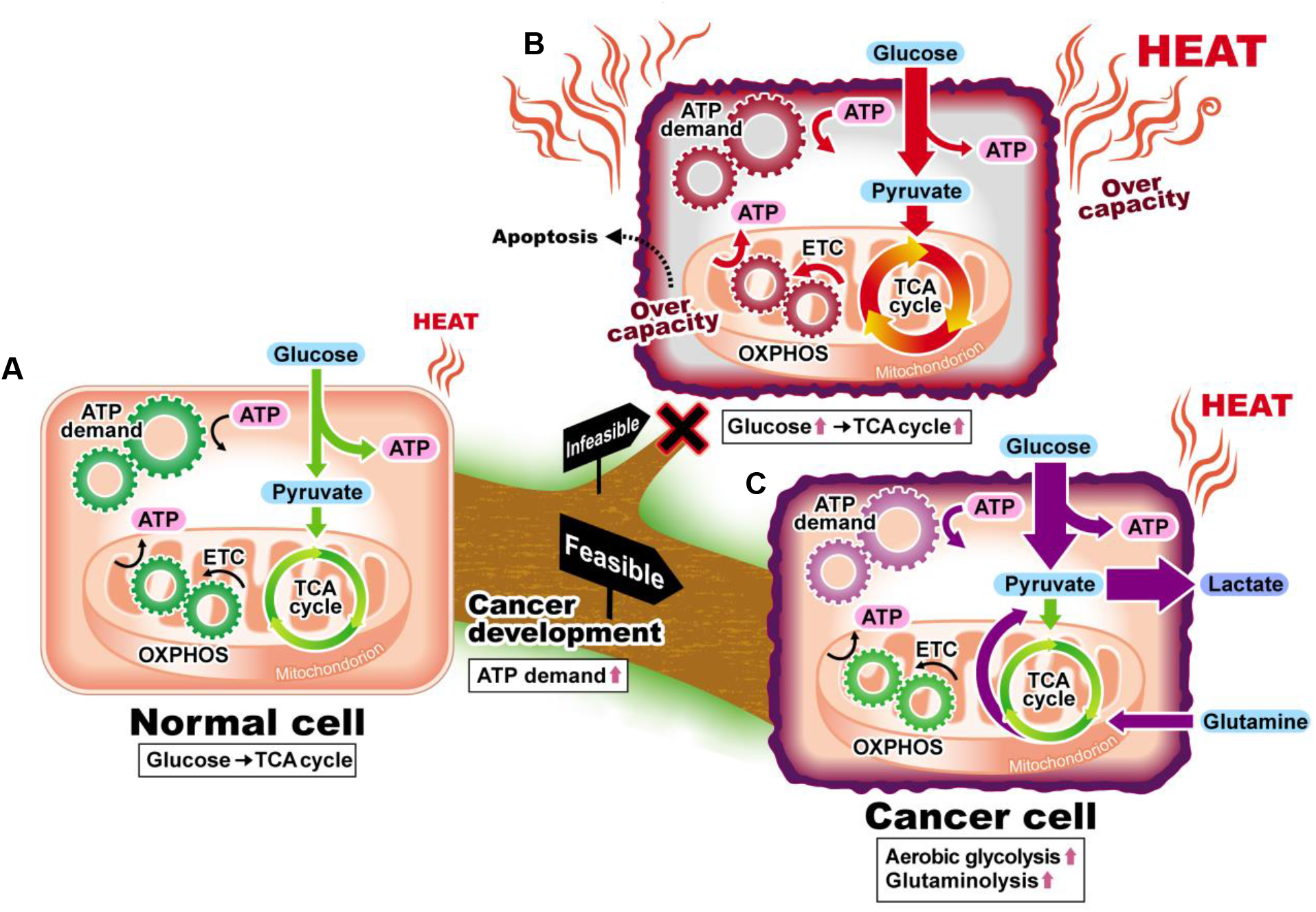
Cancer cell metabolism provides a large supply of ATP with less heat generation. **(A)** Normal cells can be assumed to regenerate ATP through the glucose → TCA cycle pathway and maintain metabolic heat dissipation and ETC activity within adequate levels. **(B)** Cancer cells have a high demand for ATP but the additional activation of the glucose → TCA cycle pathway is limited as the ETC activity and metabolic heat dissipation would otherwise exceed the permissible range. **(C)** To avoid this scenario, cancer cells rewire their metabolism to favor aerobic glycolysis and glutaminolysis to increase their ATP supply and fulfill ETC activities, respectively, with less metabolic thermogenesis.

To meet the elevated ATP demand, the active use of the OXPHOS pathway is infeasible because of the production of excessive metabolic heat (**Fig. 5B**). Notably, intracellular heat homeostasis is maintained by a balance between the generation and dissipation of a large amount of heat. ^13^C-MFA and metabolic simulation results indicated that the amount of heat dissipated from MCF-7 cells was estimated to be on the order of 100 mJ (10^6^ cells)^-1^ h^-1^ (**Fig. 2D**). This level is substantially higher than that required to increase the temperature of cultured cells by 1 ℃ [7.4 micro J (10^6^ cells)^-1^], assuming that the cells contain 1.76 pico L of water.

In contrast to the activation of the OXPHOS pathway, the additional use of aerobic glycolysis can compensate for the lack of ATP, which limits cellular thermogenesis in the glucose → TCA cycle (**Fig. 5C**). This is the only feasible scenario for ATP production in cancer cells as all 12 cancer cell lines investigated in this study employed aerobic glycolysis in combination with OXPHOS. The relationship between heat and metabolism was elucidated by *in silico* simulation (**Fig. 2**) and validated by culture experiments at low and high temperatures (**Fig 4**). The inhibition of ETC activity was compensated by metabolic rewiring for increased aerobic glycolysis while maintaining the heat dissipation level (**Fig. 3**).

Thus, the cell metabolism operates within an appropriate range of metabolic heat generation. This indicates that external heat radiation regulates respiration, as demonstrated by hyperthermia^23^. These results provide new insights into the relevance of cancer metabolism and malignancy, cancer microenvironment interactions, and drug resistance ^6^. Heat generation may also be involved in activating immune cells and controlling the cancer microenvironment ^25, 26^. Furthermore, the relationship between metabolism and heat is universal. Notably, aerobic glycolysis has been observed in normal cells, such as astrocytes, embryos, and stem cells ^27–29^. These tissues and cells commonly require large amounts of ATP for biochemical processes; however, an embryo without blood vessels has a limited capacity to release heat. The role of aerobic glycolysis in cancer and healthy cells while maintaining heat homeostasis should be analyzed.

## Materials and Methods

### Cell lines and culture conditions

Twelve human cancer cell lines derived from colorectal cancer (HCT116, DLD-1, LS174T, and WiDr), pancreatic cancer (MIA Paca-2 and PANC-1), hepatoblastoma (HepG2), hepatocellular carcinoma (huH-1), breast cancer (MCF-7), renal cancer (Caki-1), glioblastoma (YKG-1), and sarcoma (HT1080) were obtained from the RIKEN Bioresource Research Center and Japanese Collection of Research Bioresources (JCRB) Cell Bank. Culture experiments were conducted as described previously ^30^. In particular, 1.0×10^6^ cells were seeded in 10 mL of Dulbecco’s modified Eagle’s medium (DMEM) supplemented with 10% fetal bovine serum (FBS) and 1% penicillin/streptomycin (Wako) in 100-mm (diameter) plates and cultured for 15 h at 37 °C under 5% CO_2_. ^13^C-labeling was performed in 10 mL of DMEM containing 20 mM [1- ^13^C] of glucose (Cambridge Isotope Laboratories, Andover, MA, USA, over 99% purity) and 10% dialyzed FBS (Life Technologies, Gaithersburg, MD, USA). The cells were counted noninvasively at 0, 12, 16, 20, and 24 h in triplicate cultures using a CKX53 inverted microscope (Olympus, Tokyo, Japan) equipped with a DP22 digital camera (Olympus) and CKX-CCSW software (Olympus). Cell images were obtained at 10 different locations in the culture dish, and the average cell numbers were used to generate the growth curve. Culture media (200 µL) were sampled at each time point. Trypsin was added to the plates and activated at 37 °C for 1 min to measure the diameter of the floating cells. After cell collection, the diameter of live cells was counted using trypan blue staining and a TC20 automated cell counter (Bio-Rad, Hercules, CA, USA). Cell sizes were analyzed using the ImageJ software (National Institutes of Health, Bethesda, MD, USA).

### Extracellular metabolite measurements

The collected culture medium was mixed with an equal volume of 20 mM pimelate solution (internal standard) and filtered through a filter cartridge (pore size of 0.45 µm). Glucose, lactate, and acetate concentrations were determined by high-performance liquid chromatography (HPLC) equipped with a refractive index detector (Prominence, Shimadzu, Kyoto, Japan) and an Aminex HPX-87H column (Bio-Rad), as previously described ^30^. Amino acid concentrations in the culture media were measured by HPLC using the AccQ•Tag method (Armenta et al., 2010). HPLC Prominence (Shimadzu) system equipped with a Luna C18 (2) column (250 mm, 4.6 mm, and 5.0 µm, SHIMADZU GLC, Kyoto, Japan) and a photodiode array detector (260 nm) was used. Derivatized amino acids were eluted with a 20 mM sodium acetate solution containing 0.04% (v/v) of trimethylamine and phosphate adjusted to a pH of 6.8 (A) and acetonitrile (B) under the following gradient conditions: 0 min, 0% (B); 0.5 min, 8% (B), 17.5 min, 12% (B), 19 min, 15% (B), 20 min, 20% (B), 30.6 min, 100% (B), and 33.1 min, 0% (B) at a flow rate of 1.0 mL/min. The column oven temperature was maintained at 40 °C.

### Extraction and derivatization of intracellular metabolites

Intracellular metabolites were extracted using the methanol/water/chloroform method ^30^. Cellular metabolites were rapidly quenched by adding 800 µL of precooled methanol after rapid medium removal and rinsing with phosphate-buffered saline (1 mL). This procedure was performed within 15 s. Cells and solutions were collected by scraping. Cell lysates were transferred into fresh sample tubes, followed by the addition of 800 µL of cold chloroform and 320 µL of cold water. After vortexing and centrifugation, 250 µL of the upper aqueous layer was collected and dried in a vacuum concentrating centrifuge (CVE-3110; Eyela, Tokyo, Japan) at room temperature. The dried metabolites were methoxyaminated and *tert*-butyldimethylsilyated for gas chromatography-mass spectrometry (GC-MS) analysis, as described previously ^31^.

### Gas chromatography/mass spectrometry analysis

Gas chromatography/mass spectrometry (GC/MS) analysis was performed using a GCMS- QP2020 instrument (Shimadzu) equipped with a DB-5MS capillary column (Agilent Technologies). GC/MS was operated under electron impact (EI) ionization at 70 eV. One microliter of the sample was injected at 250 °C using helium as the carrier gas at a flow rate of 1 ml/min. To analyze central metabolite derivatives, the GC oven temperature was maintained at 60 °C, and then increased to 325 °C at 10 °C /min for a total run time of approximately 30 min. The MS source and quadrupole were maintained at 230 and 150 °C, respectively. The effects of naturally occurring isotopes were corrected ^32^.

### ^13^C-metabolic flux and mass balance analyses

The specific uptake and secretion rates of extracellular metabolites were calculated as previously described ^33^. The metabolic flux distribution was estimated by minimizing the variance-weighted residual sum of squares of measured and estimated mass isotopomer distributions of intracellular metabolites and effluxes using mfapy ^34^. The net ATP regeneration flux was calculated using flux distributions and the following assumptions: the reactions catalyzed by hexokinase, phosphofructokinase, pyruvate carboxylase, acetyl-coenzyme A citrate lyase/phosphoglycerate kinase, pyruvate kinase, and succinyl-coenzyme A lyase were assumed to be ATP consuming/regenerating reactions, respectively. NADPH is also assumed to be regenerated in reactions involving glucose 6-phosphate dehydrogenase, 6-phosphogluconate dehydrogenase, isocitrate dehydrogenase, and malate NADP^+^ oxidoreductase, and NADPH is consumed in the fatty acid and proline biosynthesis pathways. We assumed that excess NADPH was converted to NADH through a transhydrogenase reaction. NADH is regenerated in reactions involving glyceraldehyde-3 phosphate dehydrogenase, pyruvate dehydrogenase, alpha-ketoglutarate dehydrogenase, malate dehydrogenase, and transhydrogenase reactions and is consumed by lactate dehydrogenase. We assumed that excess NADH was converted to ATP through OXPHOS. The P/O ratio is assumed to be 2.5. Although FADH_2_ was also converted to ATP during OXPHOS, the P/O ratio was assumed to be 1.5. The ATP required for cell biomass synthesis was assumed to be 35 mmol/g of the dry cell weight ^17^.

### Clustering analysis

The gene expression dataset (E-MTAB-2706) of cancer cell lines was obtained from ArrayExpress (https://www.ebi.ac.uk/arrayexpress/) ^7^. Genes related to central metabolism were obtained from the KEGG pathway database (glycolysis/gluconeogenesis, pentose phosphate pathway, citrate cycle, D-glutamine and D-glutamate metabolism, and related transporters). The 62 genes were selected because the dataset did not contain missing data. The transcript per million (TPM)-normalized, log_2_-transformed, and Z-scored gene expression datasets of 622 cancer cell lines were used for hierarchical clustering using the Seaborn clustermap of Python 3.8 (Ward method combined with Euclidean distance).

### Flux balance analysis

A human genome-scale model (RECON2) ^17^ was used with the following modifications: i) 41 reactions responsible for the degradation of essential amino acids were removed from the model; ii) terms of standard enthalpy of formation (MegaJ mol^-1^) were added to intra/extracellular transport reactions (**Tables S7 and S8**); and iii) the reaction (R_ent) was added to the sum of the standard enthalpy of formation. The metabolic heat dissipation (h_out_) was calculated using the following equation:

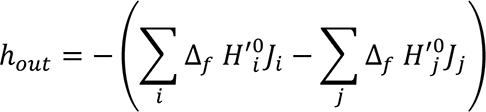

Here, *Ji* and *Jj* indicate the uptake and excretion flux levels of the *i*th and *j*th metabolite, respectively. Moreover, Δ_f_*H*^’0^ is the standard enthalpy of formation of each metabolite; Δ_f_*H*^’0^ values at a pH of 7.0 and ion strength of 0.1 M were obtained from published literature ^22^. The Δ_f_*H*^’0^ level of biomass was approximated based on the data from yeast ^35^. The approximation did not affect the FBA in this study because the metabolic flux levels for biomass synthesis were fixed at the measured values. FBA was performed using an in-house Python script with the GNU Linear Programming Kit (GLPK) in the pulp module as a linear programming solver.

### RNA-sequencing

The total RNA was extracted using the QIAzol lysis reagent (QIAGEN). Library preparation was performed using the TruSeq stranded mRNA sample prep kit (Illumina, San Diego, CA, USA) according to the manufacturer’s instructions. Sequencing was performed on an Illumina HiSeq 3000 platform in the 101 bp single-end mode. Illumina Casava1.8.2 software was used for base calling. Sequenced reads were mapped to the human reference genome sequences (hg19) using TopHat v2.0.13 in combination with Bowtie2 ver. 2.2.3 and SAMtools ver. 0.1.19. Fragments per kilobase of exon per million mapped fragments (FPKMs) were calculated using Cufflinks version 2.2.1. Gene enrichment analysis was conducted using the Enrichr web tool ^24^.

### Fluorescence microscopy

Cells were seeded on glass bottom plates (35 mm diameter, Matsunami Glass, Osaka, Japan) and stained according to the manufacturer’s protocol. The mitochondria were stained using MitoBright LT Red (Dojindo, Kumamoto, Japan). The mitochondrial membrane potential was detected using the JC-1 MitoMP Detection Kit (Dojindo). ECLIPSE TE2000-E inverted microscope (Nikon, Tokyo, Japan) equipped with an oil immersion objective lens (Plan Apo 60×/1.4 Oil Ph3 DM, Nicon), filter sets (excitation filter, 465-496 nm; emission filter, 515-555; dichroic mirror, 505, Nicon), and an iXon EMCCD camera (Andor Technology Ltd, Belfast, UK) were used. The intracellular temperature was measured using a cellular thermoprobe to determine the fluorescence ratio (Funakoshi, Tokyo, Japan). IX71 inverted microscope (Olympus, Tokyo, Japan) equipped with an objective lens (LCPlanFl 20×/0.40 Ph1, Olympus) and ORCA-Spark digital CMOS camera (Hamamatsu) were used. Fluorescence images were taken using U-MWIBA3 (excitation filter, 460-495 nm; emission filter, 510-550 nm; dichroic mirror, 505 nm, Olympus) and custom-made (excitation filter, 460-495 nm; emission filter, 570-625 nm; dichroic mirror, 505 nm, Olympus) filter cubes. The temperature and CO_2_ content of cells were maintained at 37 ℃ and 5%, respectively, using a microscope incubator (BLAST, Kawasaki, Japan). Images were analyzed using the ImageJ software.

## Acknowledgments

We acknowledge the NGS core facility of the Genome Information Research Center at the Research Institute for Microbial Diseases, Osaka University for their support in RNA sequencing and data analysis. We thank Prof. Shinya Kuroda, Dr. Satoshi Ohno (University of Tokyo), and Shimpei Kawaoka (Kyoto University) for helpful comments on this manuscript. We also thank Keiko Fukamoto for her skillful technical support. Grant-in-Aid for Scientific Research on Innovative Areas (Grant no. 17H06303). Extramural Collaborative Research Grant of Cancer Research Institute, Kanazawa University (Grant no.04-13).

## Author contributions

N.O., S.K., H.S., C.T. and F.M. designed the study. N.O., T.S., Y.K., C.A., and H.U. performed ^13^C-tracer experiments. F.M. performed flux balance analysis. N.O., S.T. and A.S. performed metabolome analysis. N.O. performed microscopic analysis and RNA-sequencing. N.O., T.S., Y.K., C.A., S.T., A.S., H.U., S.K., H.S., C.T. and F.M. analyzed and interpreted data. N.O. and F.M. wrote original draft. N.O., H.S., S.K., C.T. and F.M. reviewed and edited manuscript with help from all authors.

## Data availability

RNA-Seq data is available at DDBJ of the National Institute of Genetics (https://www.ddbj.nig.ac.jp/index-e.html) via the index of DRA012975

## Supplementary Materials

### Supplementary Text

#### Text S1. Detailed explanation of the ^13^C-metabolic flux analysis procedure

The metabolic model consisted of the major pathways of central carbon metabolism, including glycolysis, the pentose phosphate pathway, the TCA cycle, anaplerotic reactions, and biomass synthesis (**Table S8**). Folate metabolism was ignored, as MTHFD1 and MTHFD2 were not highly expressed (**Fig. S1**). The catabolism of the essential amino acids was ignored. Biomass effluxes of amino acids for proteins, glucose-6-phosphate for glycogen, ribose-5- phosphate for ribonucleic acid, and cytosolic acetyl-CoA for fatty acids were calculated from the specific growth rates and biomass compositions of mammalian cells ^36^. The constant for the conversion from cell number to dry cell weight was assumed to be 0.392 mg-dry cell weight (10^6^ cells)^-1^ for all cell lines ^36^. This assumption was based on the observation that the detached cell sizes from culture dishes were almost identical to all cell lines except for Caki-1 (**Table S1**). Additionally, the effect of this assumption on the total metabolic flux was considered to be minor because the effluxes for biomass synthesis accounted for only a minor percentage of carbon uptake flux. The goodness of fit of ^13^C-MFA was evaluated by χ^2^ statistics test, where the standard deviations of ^13^C-labeling were assumed to be 0.01 ^31^. The ^13^C-MFA of all cultured cancer cell lines successfully passed the χ^2^-test (**Table S2**). All 95% confidence intervals were determined using the grid search method and are shown in **Table S2** ^37^.

#### Text S2. Comparison of the net ATP regeneration rate in cultured cancer cells with that of normal cells

In this study, the net ATP regeneration rate of MCF-7 cells was determined as 1815 nmol (10^6^ cells)^-1^ h^-1^ based on the metabolic flux distribution (**Fig. 1a**). The net ATP regeneration rates ranged from 1460 (LS174T) to 4086 (Caki-1) nmol (10^6^ cells)^-1^ h^-1^ among the 12 cell lines. The total human cell number is 3.72 × 10^13^, composed of 15% non-erythrocyte cells ^38^. Moreover, ATP was presumed to be regenerated by OXPHOS using glucose (32 ATP/glucose). If all non-erythrocyte cells reproduce ATP at the same level of LS174T, the required amount of glucose (g day^-1^) is 1460×10^-9^ nmol (10^-6^ cells)^-1^ h^-1^×24 h×3.72×10^13^ cells×0.15×1/32 (glucose/ATP) ×180 g mol^-1^ = 1100 g day^-1^, equal to 3686 kcal day^-1^. The recommended daily calorie intake is 2,000 and 2,500 kcal day^-1^ for women and men, respectively, suggesting that the ATP regeneration level of LS174T cells was higher than that of normal cells.

Moreover, the metabolic flux analysis conducted in this study revealed that the ATP regeneration rates by OXPHOS in 12 cancer cells were 447–1727 nmol (10^6^ cells)^-1^ h^-1^. If the glucose equivalent of 2,500 kcal day^-1^ were catabolized to produce ATP by OXPHOS, the net ATP regeneration rate would be 994 nmol (10^6^ cells)^-1^ h^-1^. This level is similar to reported data such as that of rat heart H9c2 (normal) cells (1282.5 nmol (10^6^ cells)^-1^ h^-1^) ^39^.

### Supplementary figures

**Figure S1.**
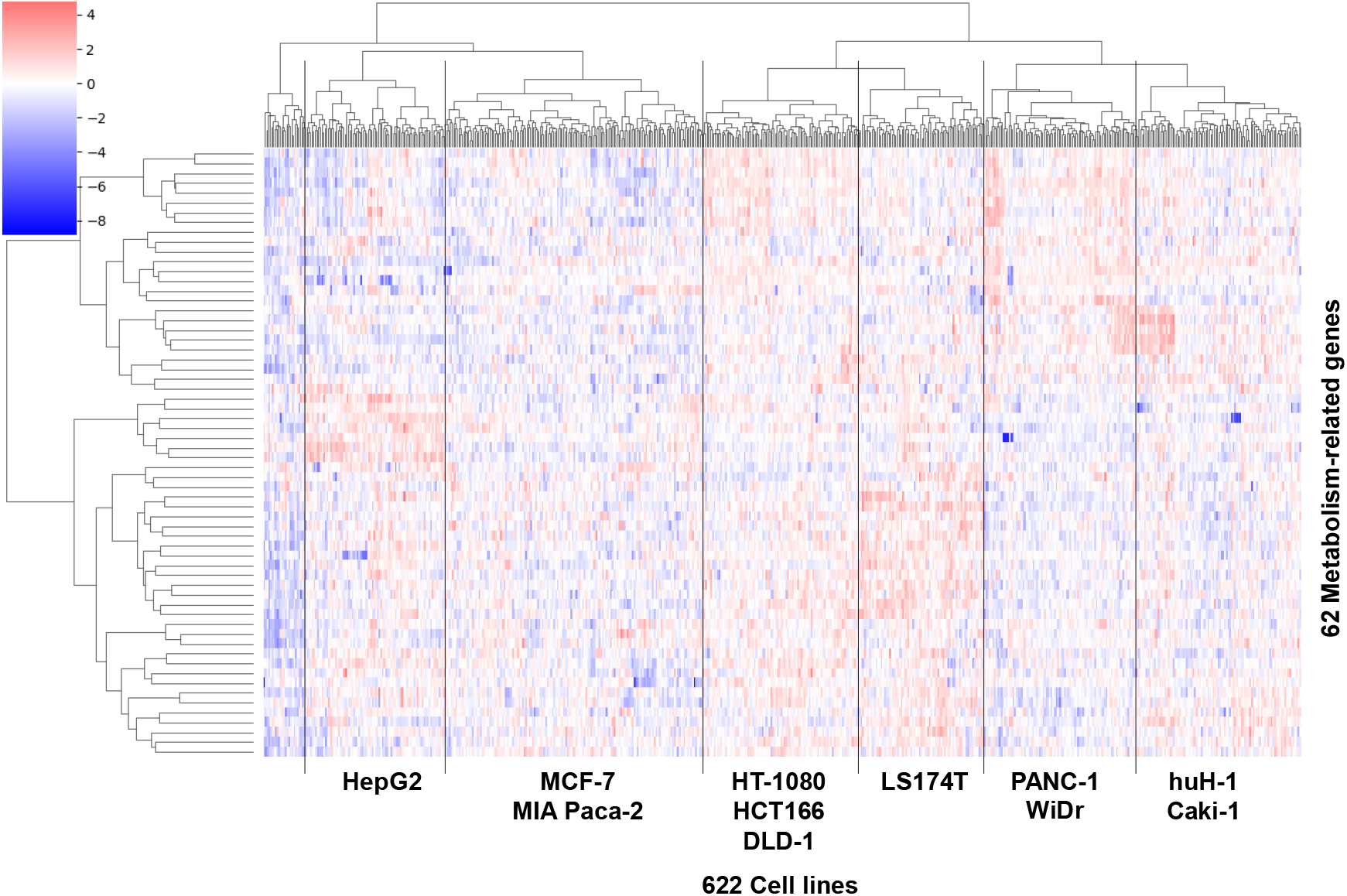
Hierarchical clustering of 622 cancer cell line using publicly available expression data of 62 central metabolism-related genes. 62 central carbon metabolism-related genes were obtained from the KEGG pathway database and selected due to there being no missing data in the dataset. For hierarchical clustering, the Ward method combined with the Euclidean distance was used. From each cluster, colorectal cancer (HCT116, DLD-1, LS174T, and WiDr), pancreatic cancer (MIA Paca-2, and PANC-1), hepatoblastoma (HepG2), hepatocellular carcinoma (huH-1), breast cancer (MCF-7), renal cancer (Caki-1), and sarcoma (HT1080) were selected. Moreover, glioblastoma (YKG-1) was arbitrarily selected for the analysis.

**Figure S2.**
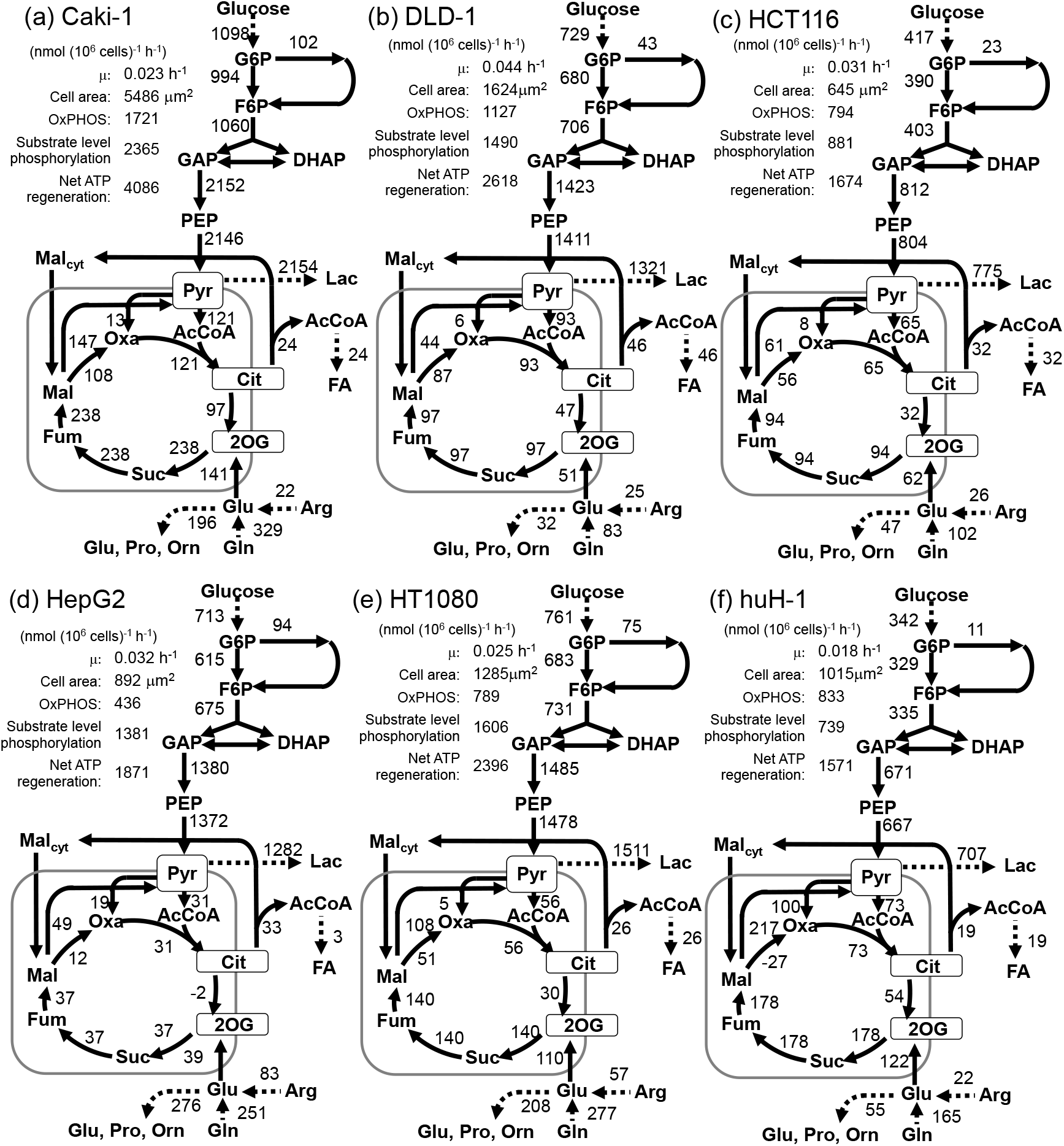
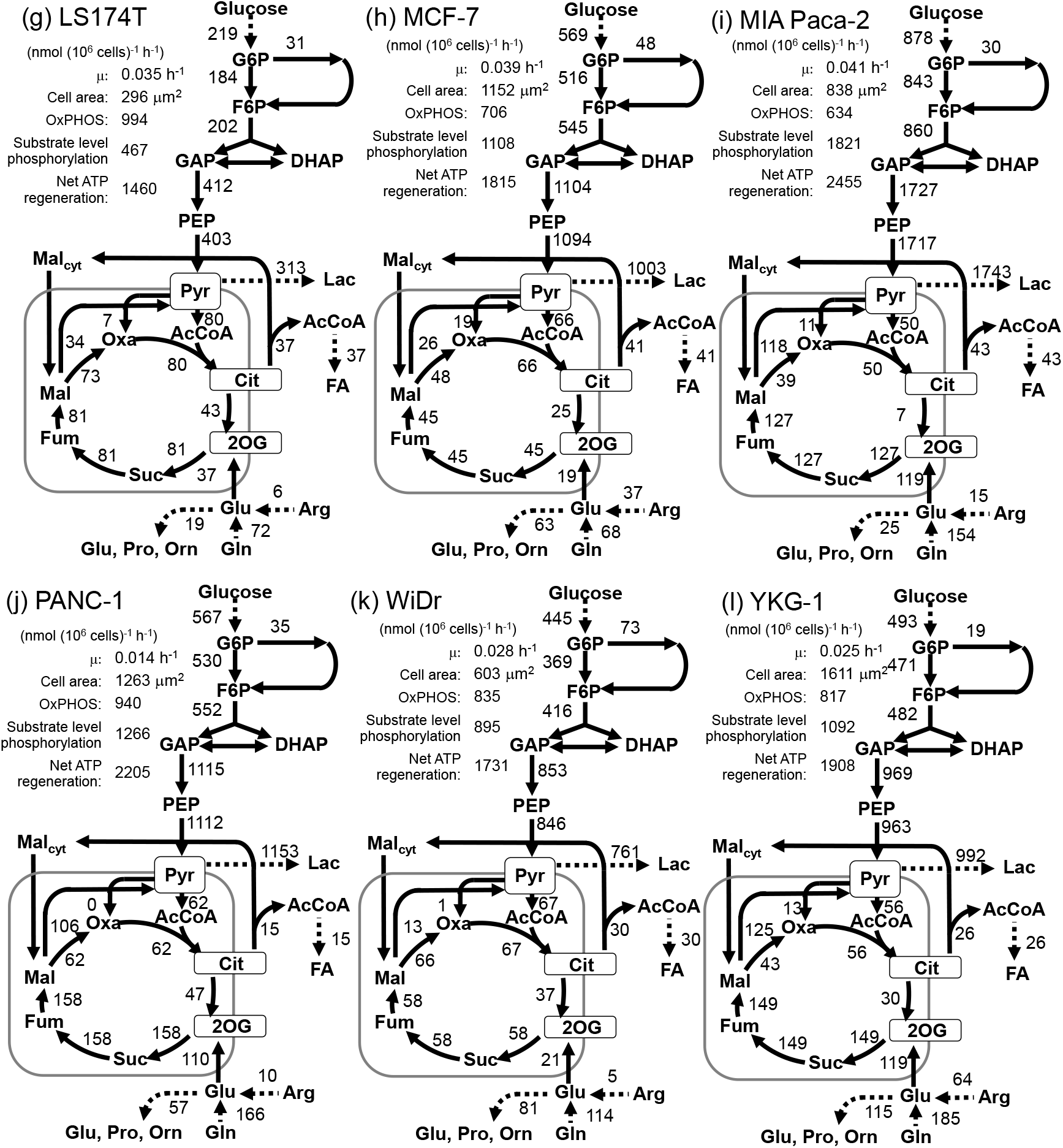
Metabolic flux distribution of 12 cancer cell lines as determined by ^13^C-MFA. All the values represent the metabolic flux level (nmol (10^6^ cells)^-1^ h^-1^). Measured specific cell proliferation rate (µ[h^-1^]) and cell adhesion area (cell area [µm^2^]), as well as the ATP regeneration rates by OXPHOS, substrate-level phosphorylation, and the net ATP regeneration rate are shown in the figure.

**Figure S3.**
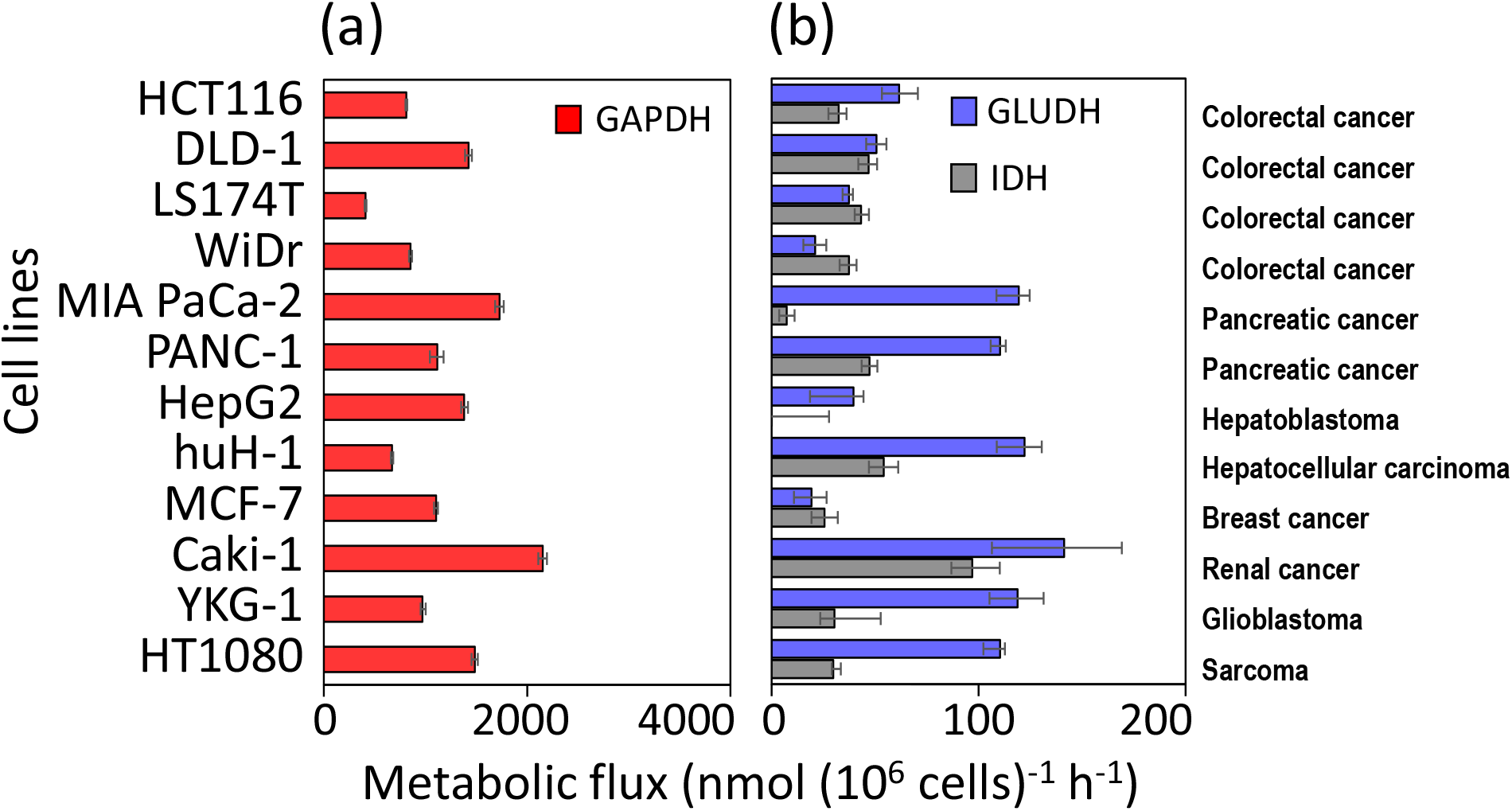
Variations in flux levels in representative metabolic pathways. Flux levels of (a) GAPDH and (b) GLUDH and IDH reactions are presented. Error bars represent 95% confidence intervals.

**Figure S4.**
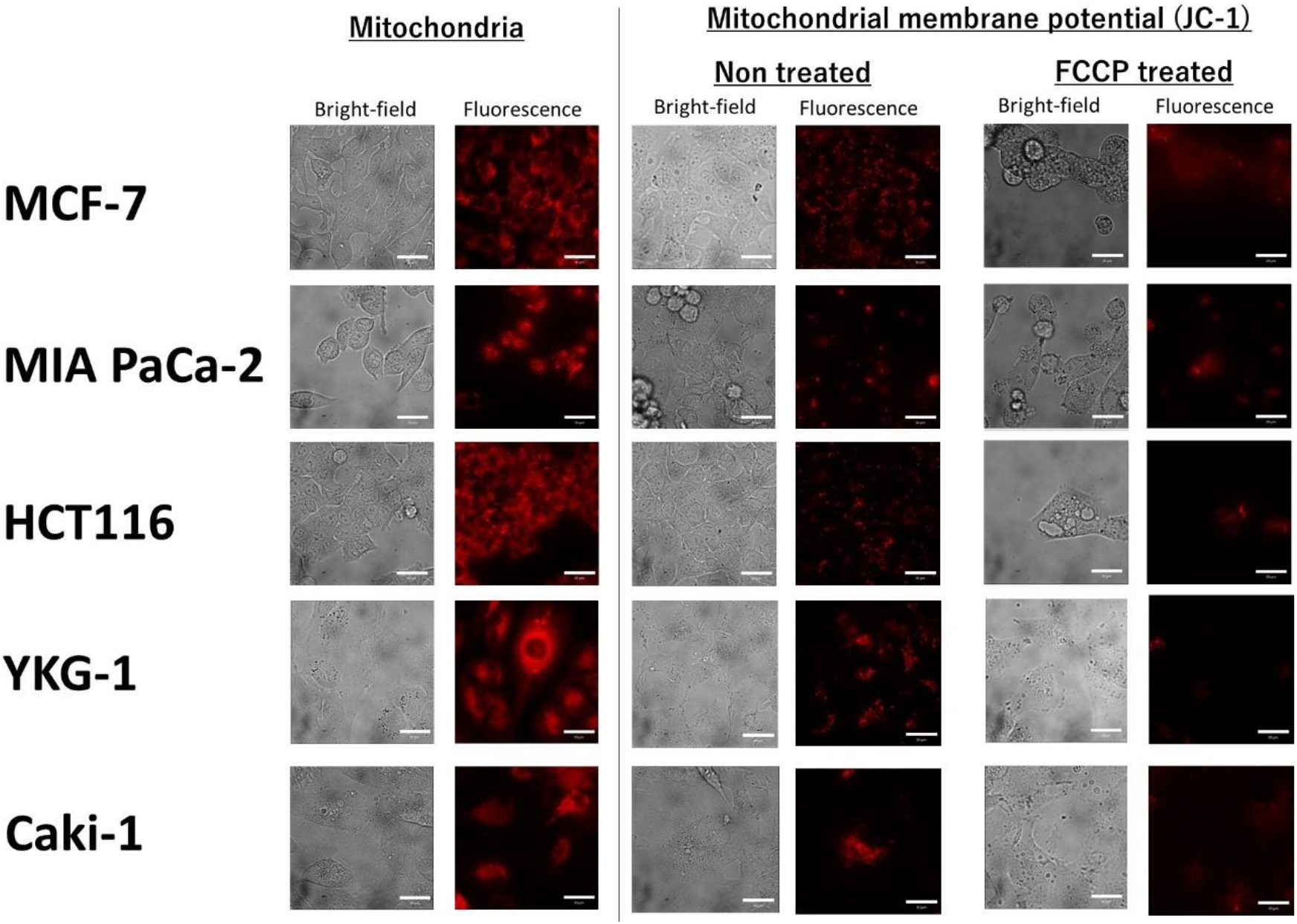
Mitochondrial membrane potential is retained in cultured cancer cells. The potential-dependent dye, JC-1, was used to monitor the mitochondrial membrane potential.

**Figure S5.**
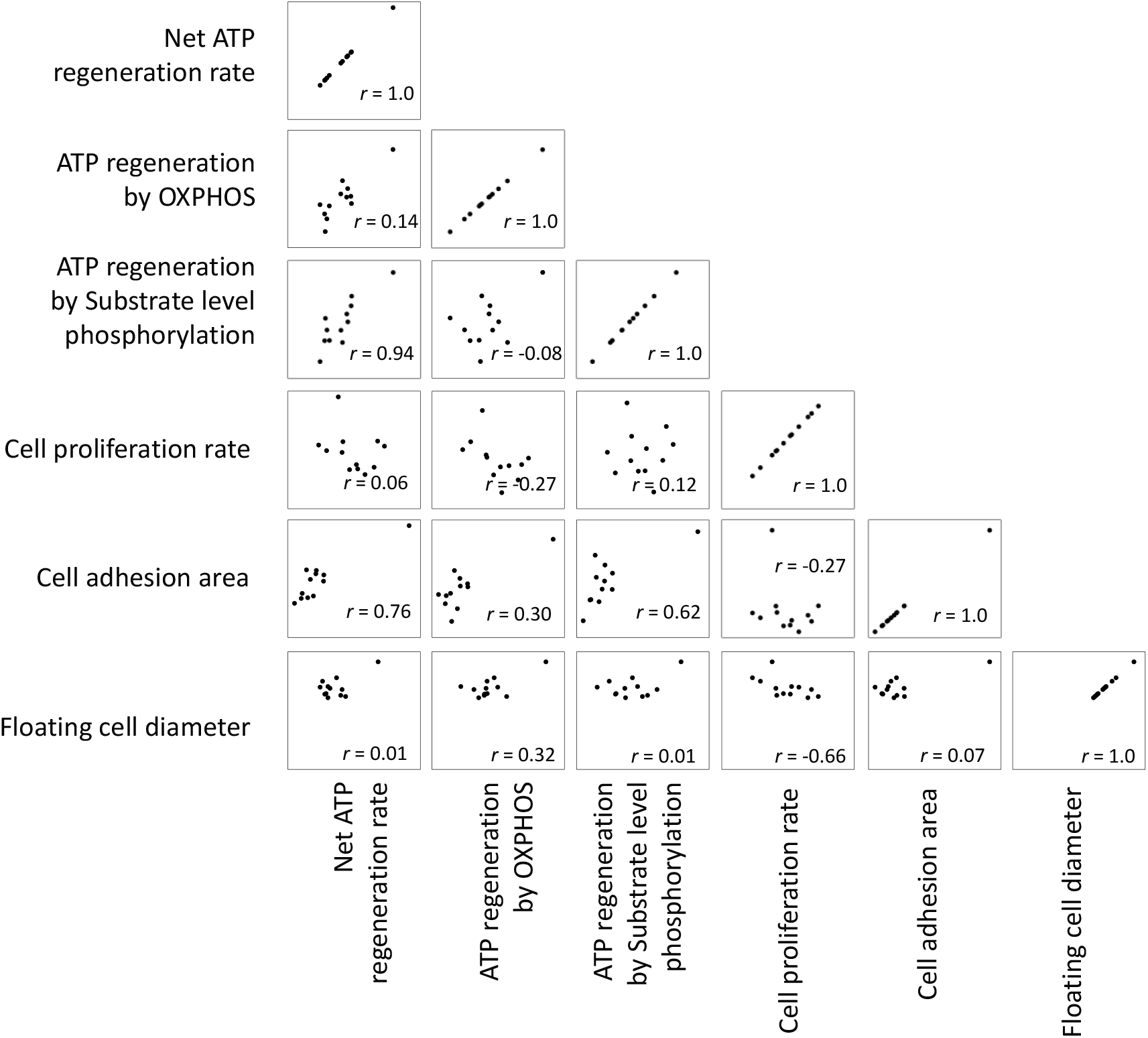
Scatter plots among the ATP regeneration rates and the visible phenotype data. Spearman’s rank order correlation coefficients were also shown.

**Figure S6.**
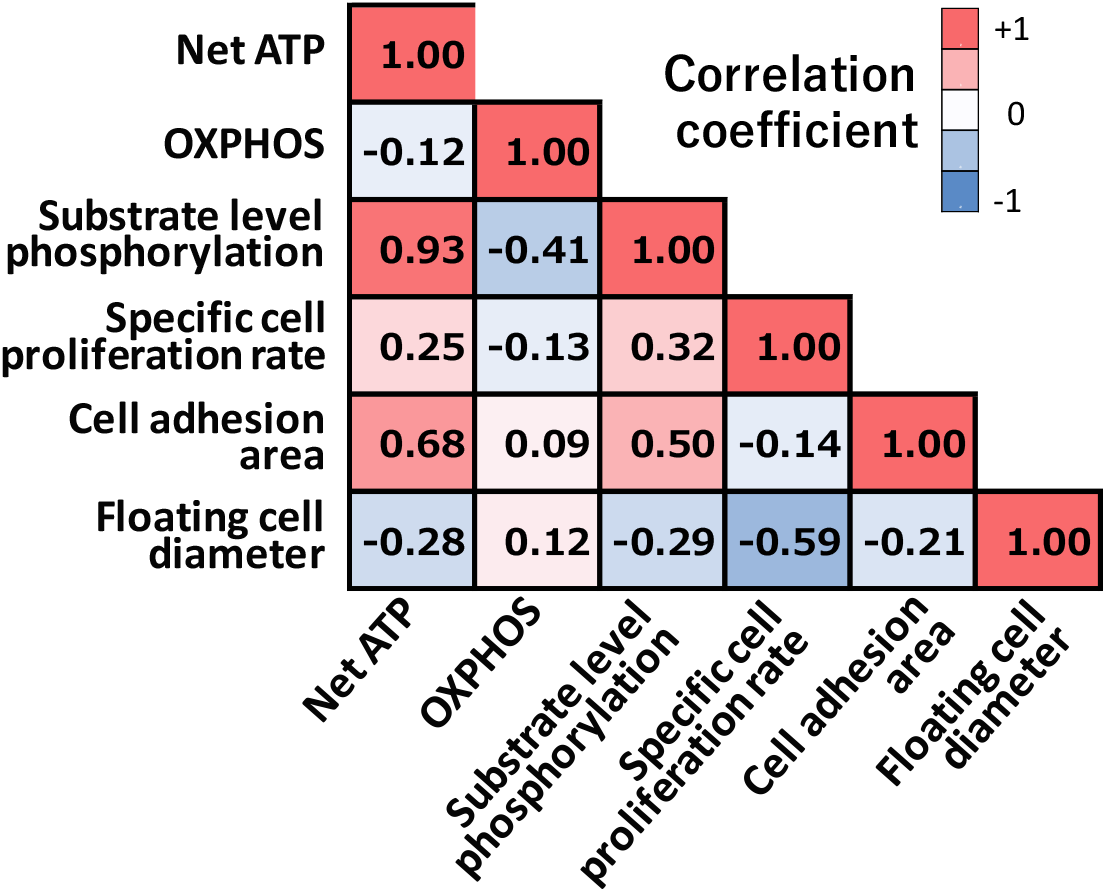
Correlation among ATP regeneration rates and the visible phenotypes in the cell upon removal of the outlier cell line (Caki-1) from the dataset. Spearman’s rank order correlation coefficients are shown in the heat map (*n*=11).

**Figure S7.**
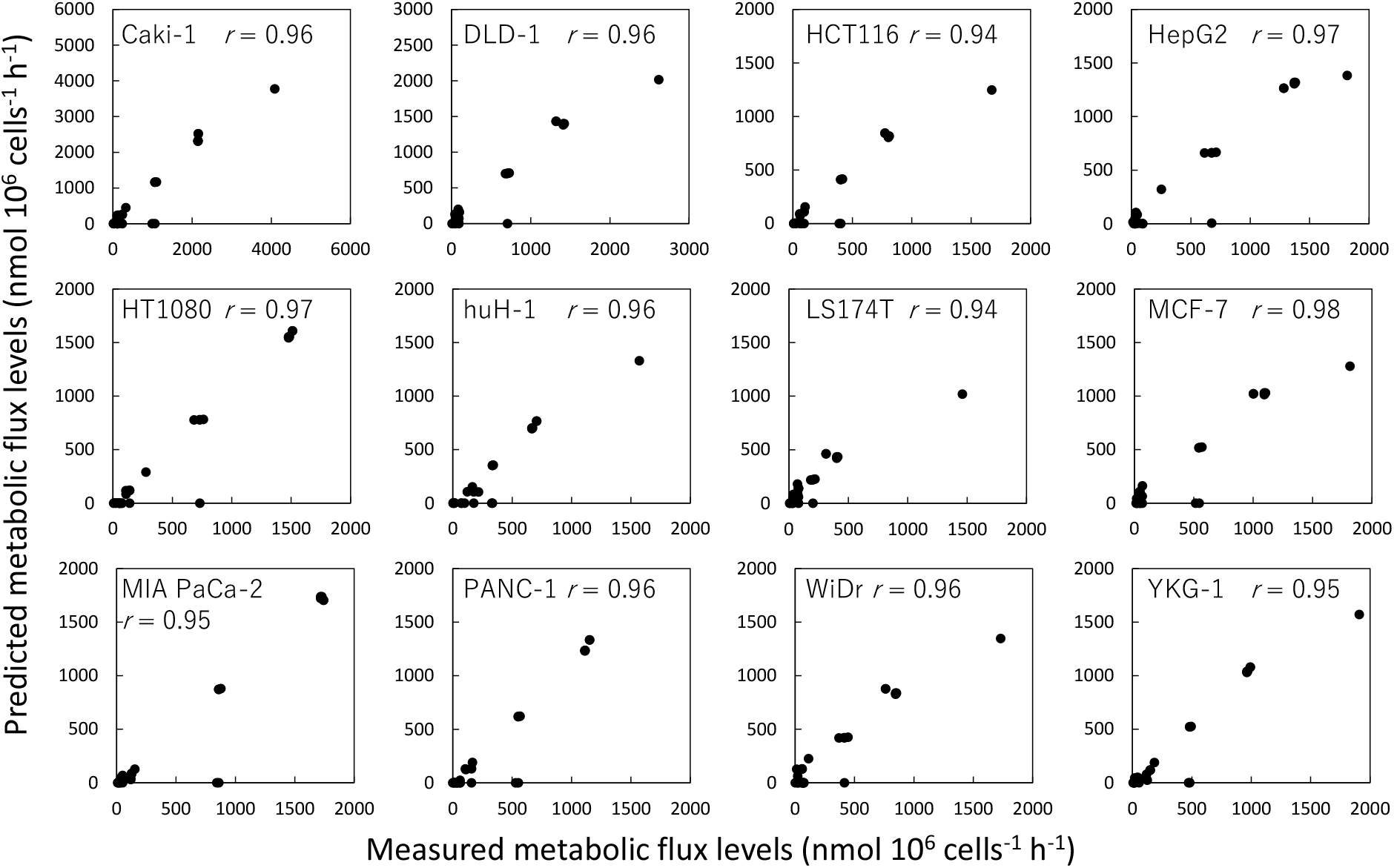
Comparison of measured metabolic flux data and predicted metabolic flux distributions by FBA for 12 cell lines. *In silico* simulation of metabolism was performed by FBA using a genome scale model of human metabolism (RECON2).

### Supplementary tables

**Table S1.** Culture profiles of 12 cell lines.

| Measured | Unit | Caki-1 | DLD-1 | HCT116 | HepG2 | HT-1080 | Huh-1 |
| --- | --- | --- | --- | --- | --- | --- | --- |
| Specific cell growth rate | $\text{h}^{-1}$ | 0.0228 | 0.0437 | 0.031 | 0.0316 | 0.0249 | 0.0175 |
| cell adhesion area | $\mu\text{m}^2$ ($n = 30$) | $3486 \pm 792$ | $1623 \pm 468$ | $645 \pm 159$ | $891 \pm 300$ | $1284 \pm 533$ | $1015 \pm 322$ |
| Diameter of floating cells | $\mu\text{m}$ ($n = 56-595$) | $18.3 \pm 3.7$ | $12.4 \pm 2.2$ | $12.8 \pm 1.5$ | $14.1 \pm 1.5$ | $12.6 \pm 1.9$ | $15.0 \pm 2.0$ |
| Glucose uptake | $\text{nmol} (10^6 \text{ cells})^{-1} \text{ h}^{-1}$ | $606 \pm 232$ | $742 \pm 32.0$ | $361 \pm 17.0$ | $755 \pm 250$ | $604 \pm 75.8$ | $397 \pm 45.0$ |
| Glutamine uptake | $\text{nmol} (10^6 \text{ cells})^{-1} \text{ h}^{-1}$ | $327 \pm 30$ | $84 \pm 4.8$ | $98 \pm 5.3$ | $295 \pm 46$ | $278 \pm 7.07$ | $168 \pm 9.3$ |
| Arginine uptake | $\text{nmol} (10^6 \text{ cells})^{-1} \text{ h}^{-1}$ | $22 \pm 2$ | $25 \pm 1.2$ | $25 \pm 1.2$ | $84 \pm 6$ | $56.9 \pm 2.78$ | $22 \pm 2.2$ |
| Valine uptake | $\text{nmol} (10^6 \text{ cells})^{-1} \text{ h}^{-1}$ | $-32.9 \pm 21.3$ | $7 \pm 0.7$ | $5 \pm 1.6$ | $66 \pm 26$ | $8.5 \pm 4.8$ | $13.6 \pm 2.8$ |
| Leucine uptake | $\text{nmol} (10^6 \text{ cells})^{-1} \text{ h}^{-1}$ | $4.2 \pm 17.2$ | $16 \pm 0.5$ | $12 \pm 1.9$ | $76 \pm 22$ | $22.2 \pm 4.9$ | $17.5 \pm 2.0$ |
| Isoleucine uptake | $\text{nmol} (10^6 \text{ cells})^{-1} \text{ h}^{-1}$ | $-24.4 \pm 22.5$ | $9.4 \pm 0.4$ | $13.9 \pm 2.7$ | $76 \pm 22$ | $17.5 \pm 5.2$ | $32.9 \pm 5.3$ |
| Lactate excretion | $\text{nmol} (10^6 \text{ cells})^{-1} \text{ h}^{-1}$ | $2158 \pm 27$ | $1311 \pm 40$ | $776 \pm 3$ | $1282 \pm 29$ | $1514 \pm 13.0$ | $708 \pm 3$ |
| Glutamate excretion | $\text{nmol} (10^6 \text{ cells})^{-1} \text{ h}^{-1}$ | $143 \pm 5$ | $1 \pm 1$ | $22 \pm 0$ | $132 \pm 14$ | $133 \pm 2.96$ | $43 \pm 2$ |
| Alanine excretion | $\text{nmol} (10^6 \text{ cells})^{-1} \text{ h}^{-1}$ | $0 \pm 0$ | $23.2 \pm 1.3$ | $8.8 \pm 0.2$ | $80 \pm 7$ | $7.14 \pm 0.317$ | $0.0 \pm 0.0$ |
| Proline excretion | $\text{nmol} (10^6 \text{ cells})^{-1} \text{ h}^{-1}$ | $34.6 \pm 0.4$ | $9.2 \pm 0.5$ | $1.7 \pm 0.1$ | $21 \pm 2$ | $12.4 \pm 0.472$ | $5.4 \pm 0.3$ |
| Ornithine excretion | $\text{nmol} (10^6 \text{ cells})^{-1} \text{ h}^{-1}$ | $18.2 \pm 3.03$ | $21.9 \pm 1.2$ | $23.9 \pm 0.7$ | $112 \pm 19$ | $62.9 \pm 2.6$ | $6.4 \pm 1$ |

| Measured | Unit | LS174T | MCF7 | MIAPACA2 | Panc-1 | WiDr | YKG-1 |
| --- | --- | --- | --- | --- | --- | --- | --- |
| Specific cell growth rate | $\text{h}^{-1}$ | 0.0349 | 0.039 | 0.0406 | 0.0139 | 0.0279 | 0.0245 |
| cell adhesion area | $\mu\text{m}^2$ ($n = 30$) | $296 \pm 93$ | $1152 \pm 372$ | $838 \pm 174$ | $1263 \pm 405$ | $603 \pm 196$ | $1611 \pm 588$ |
| Diameter of floating cells | $\mu\text{m}$ ($n = 56-595$) | $14.0 \pm 2.0$ | $12.2 \pm 1.8$ | $13.6 \pm 1.8$ | $15.6 \pm 2.1$ | $12.9 \pm 1.6$ | $13.8 \pm 1.4$ |
| Glucose uptake | $\text{nmol} (10^6 \text{ cells})^{-1} \text{ h}^{-1}$ | $222 \pm 21.0$ | $507 \pm 9.9$ | $1031 \pm 81.7$ | $538 \pm 133.5$ | $481 \pm 37.0$ | $400 \pm 43.98$ |
| Glutamine uptake | $\text{nmol} (10^6 \text{ cells})^{-1} \text{ h}^{-1}$ | $75 \pm 2.7$ | $66 \pm 5.0$ | $156 \pm 7.1$ | $167 \pm 1.7$ | $114 \pm 3.8$ | $184.2 \pm 8.04$ |
| Arginine uptake | $\text{nmol} (10^6 \text{ cells})^{-1} \text{ h}^{-1}$ | $6 \pm 0.2$ | $37 \pm 1.0$ | $15 \pm 1.2$ | $9 \pm 0.5$ | $4 \pm 0.6$ | $63.76 \pm 1.26$ |
| Valine uptake | $\text{nmol} (10^6 \text{ cells})^{-1} \text{ h}^{-1}$ | $-2 \pm 1.3$ | $2 \pm 2.9$ | $17.8 \pm 2.4$ | $7.09 \pm 0.5$ | $-1 \pm 0.8$ | $24.6 \pm 3.9$ |
| Leucine uptake | $\text{nmol} (10^6 \text{ cells})^{-1} \text{ h}^{-1}$ | $9 \pm 0.5$ | $13 \pm 2.7$ | $28.2 \pm 2.7$ | $17.4 \pm 0.7$ | $4 \pm 0.8$ | $36.5 \pm 4.1$ |
| Isoleucine uptake | $\text{nmol} (10^6 \text{ cells})^{-1} \text{ h}^{-1}$ | $7.4 \pm 0.5$ | $6.0 \pm 1.9$ | $28.4 \pm 2.8$ | $15.5 \pm 1.2$ | $-0.6 \pm 0.3$ | $33.2 \pm 4.1$ |
| Lactate excretion | $\text{nmol} (10^6 \text{ cells})^{-1} \text{ h}^{-1}$ | $313 \pm 7$ | $1014 \pm 6$ | $1734 \pm 29$ | $1146 \pm 41$ | $762 \pm 10$ | $993 \pm 7.18$ |
| Glutamate excretion | $\text{nmol} (10^6 \text{ cells})^{-1} \text{ h}^{-1}$ | $13 \pm 1$ | $16 \pm 1$ | $18 \pm 1$ | $41 \pm 2$ | $53 \pm 2$ | $43.7 \pm 0.51$ |
| Alanine excretion | $\text{nmol} (10^6 \text{ cells})^{-1} \text{ h}^{-1}$ | $27.0 \pm 0.6$ | $22.0 \pm 0.8$ | $20.9 \pm 0.9$ | $0.0 \pm 0.0$ | $22.8 \pm 0.2$ | $19.75 \pm 1.11$ |
| Proline excretion | $\text{nmol} (10^6 \text{ cells})^{-1} \text{ h}^{-1}$ | $3.5 \pm 0.2$ | $4.0 \pm 0.2$ | $3.1 \pm 0.3$ | $12.1 \pm 0.1$ | $4.4 \pm 0.1$ | $11.83 \pm 0.286$ |
| Ornithine excretion | $\text{nmol} (10^6 \text{ cells})^{-1} \text{ h}^{-1}$ | $2.9 \pm 0.1$ | $43.0 \pm 1.4$ | $4.0 \pm 1.2$ | $4.69 \pm 0.1$ | $23.4 \pm 0.4$ | $59.7 \pm 1.5$ |

**Table S2.**
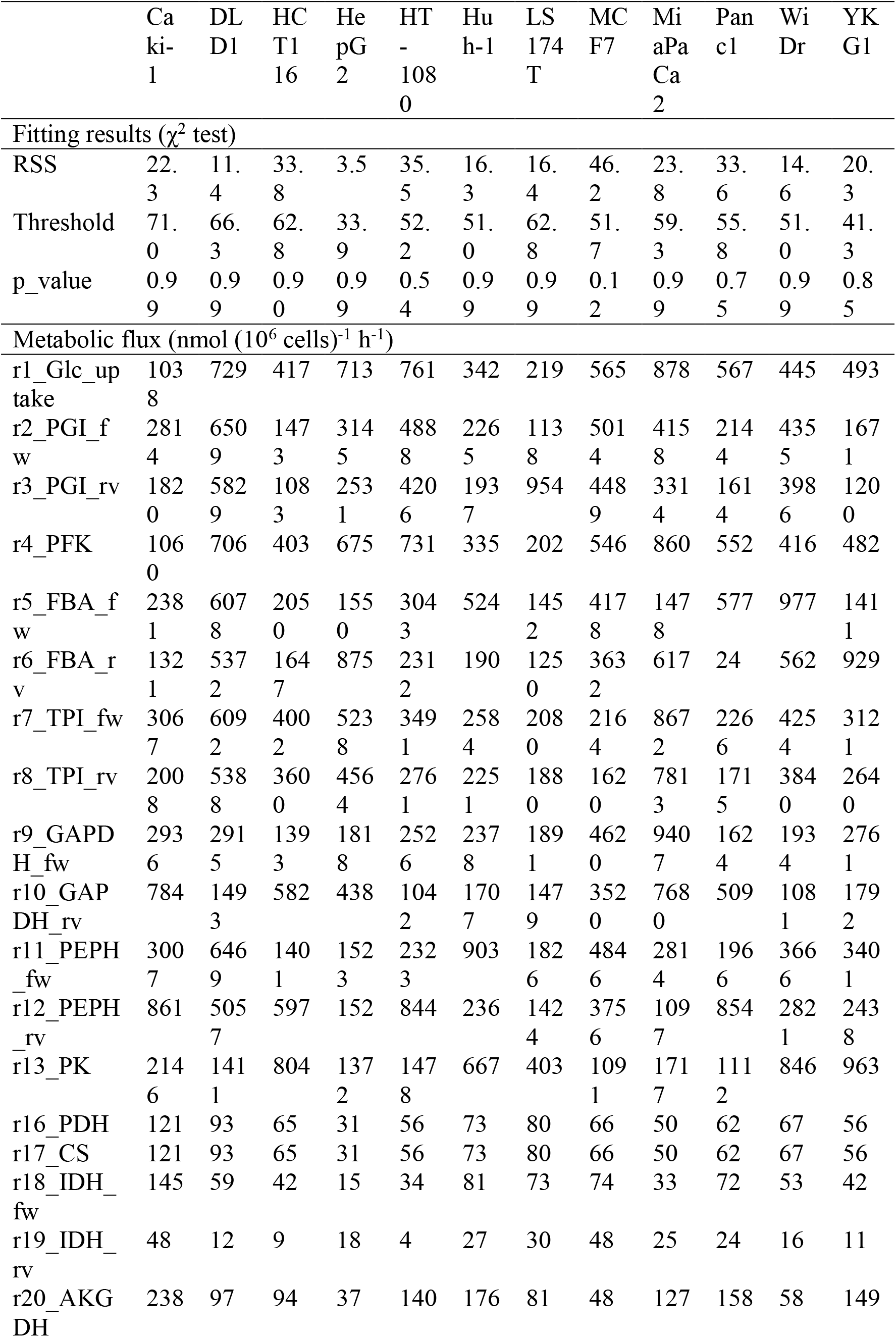
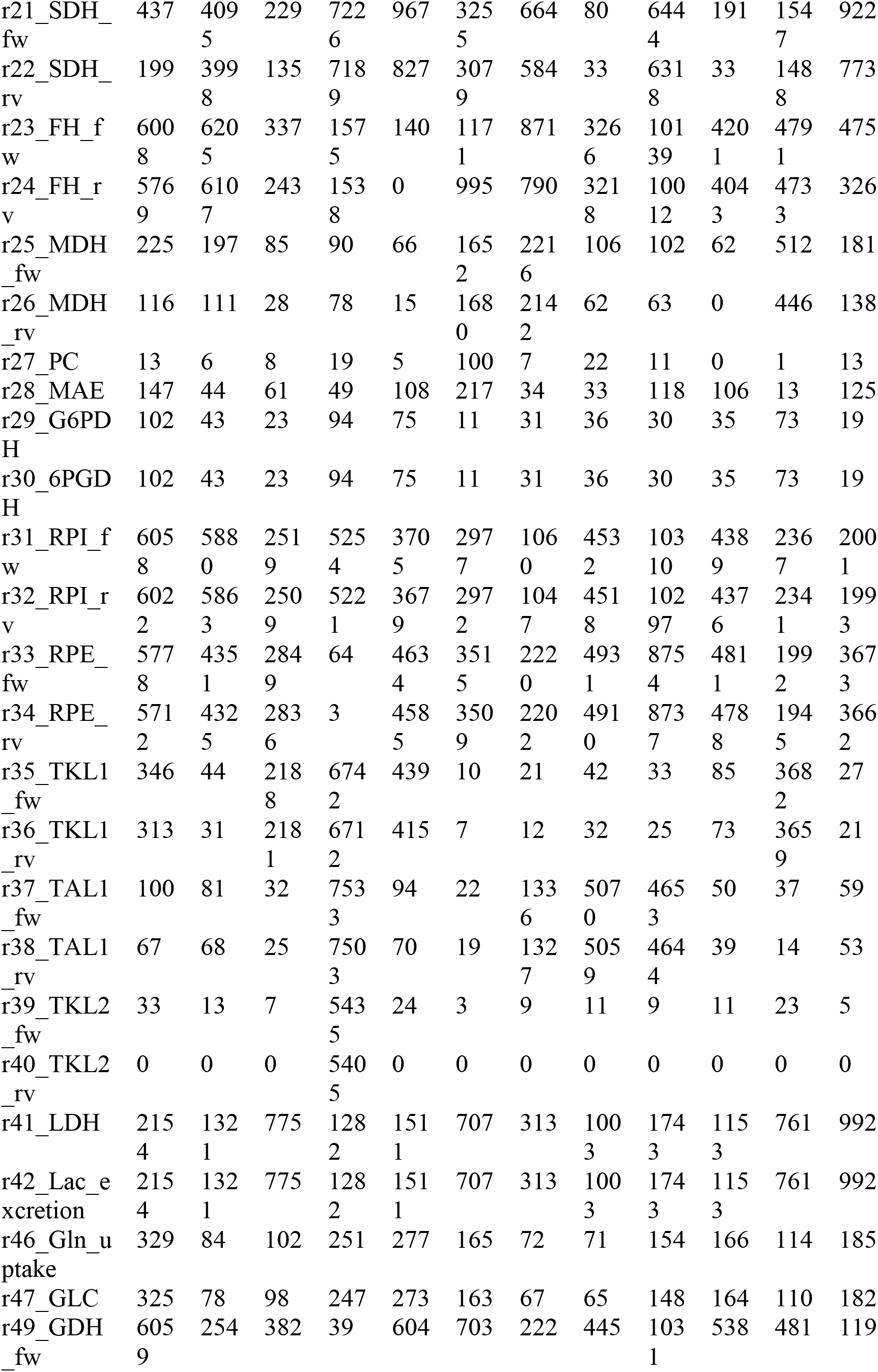
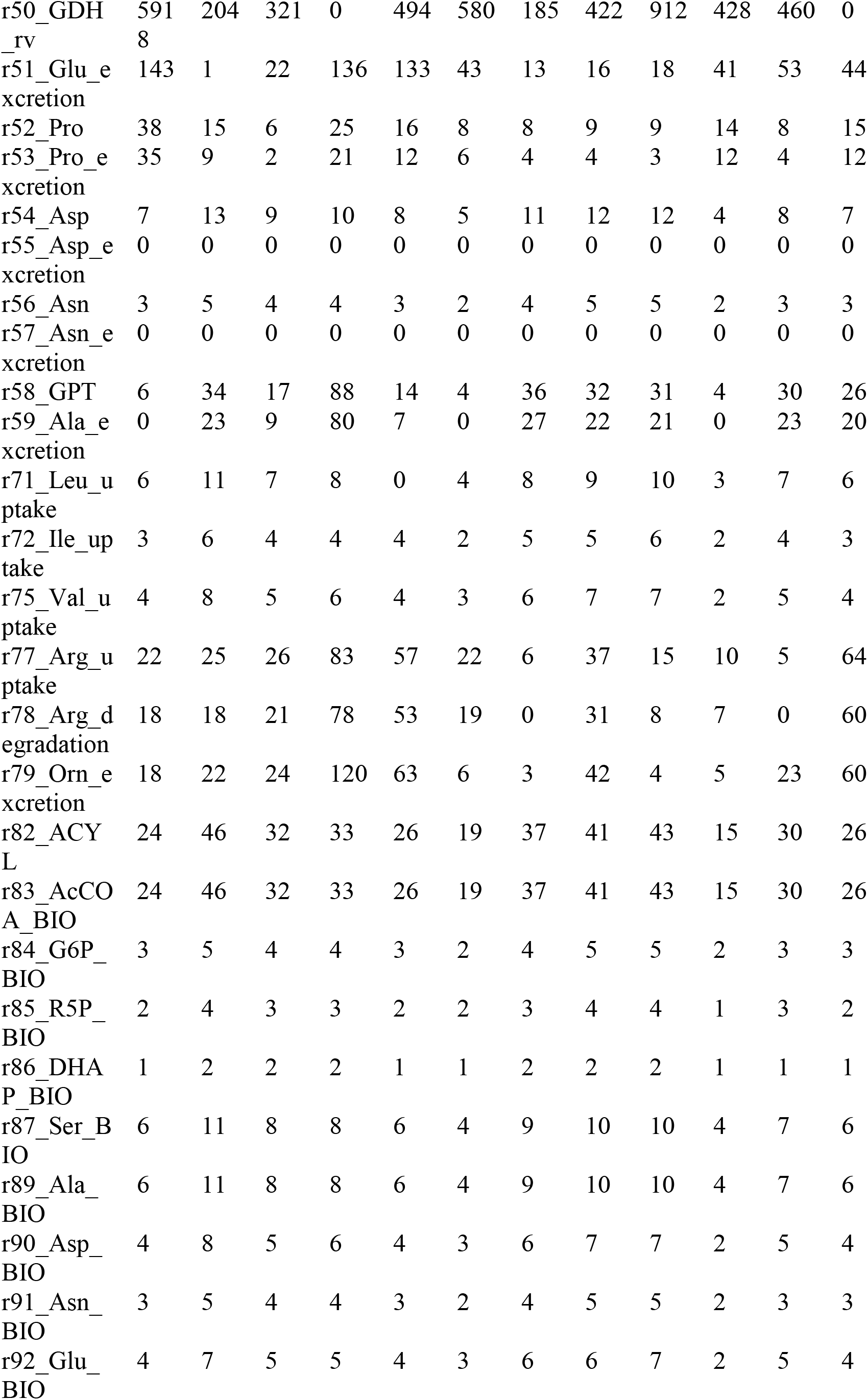
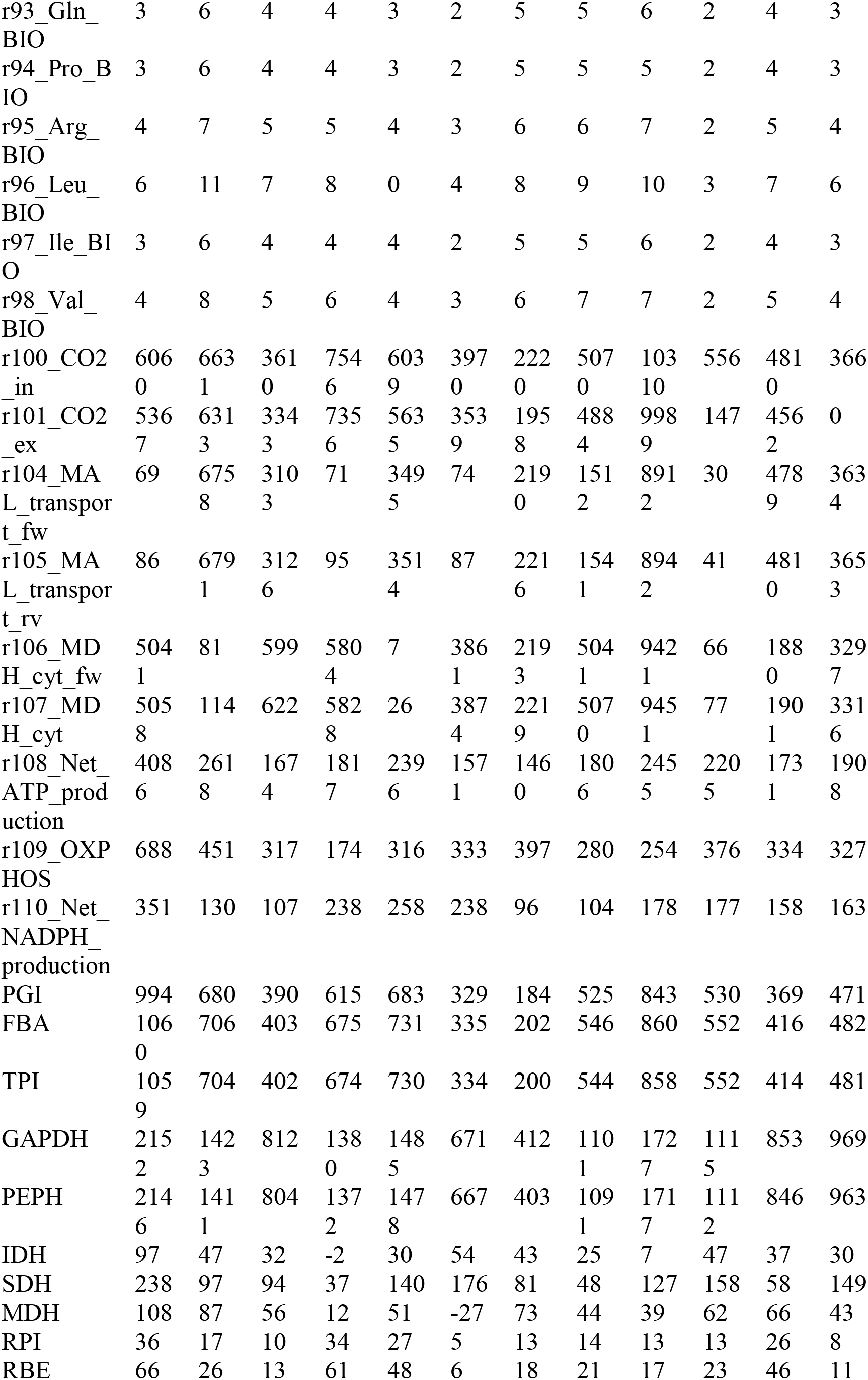
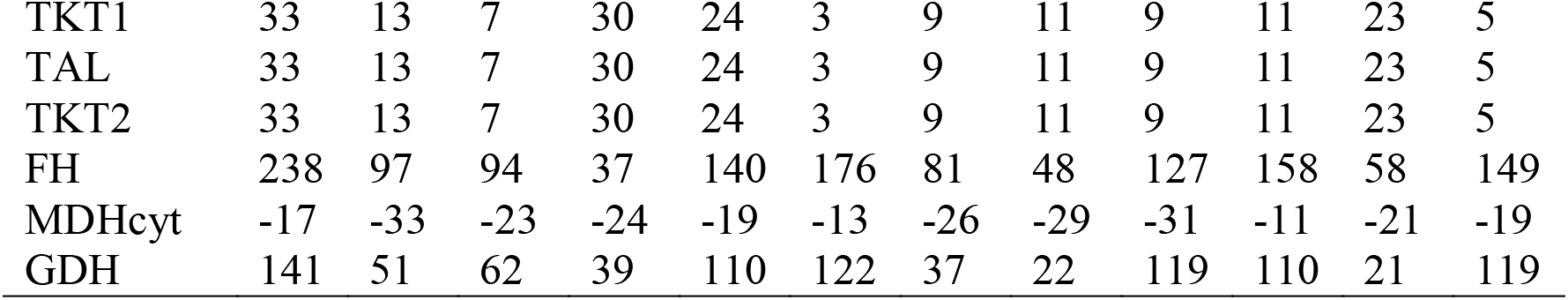
Results of ^13^C-MFA of 12 cell lines.

**Table S3.**
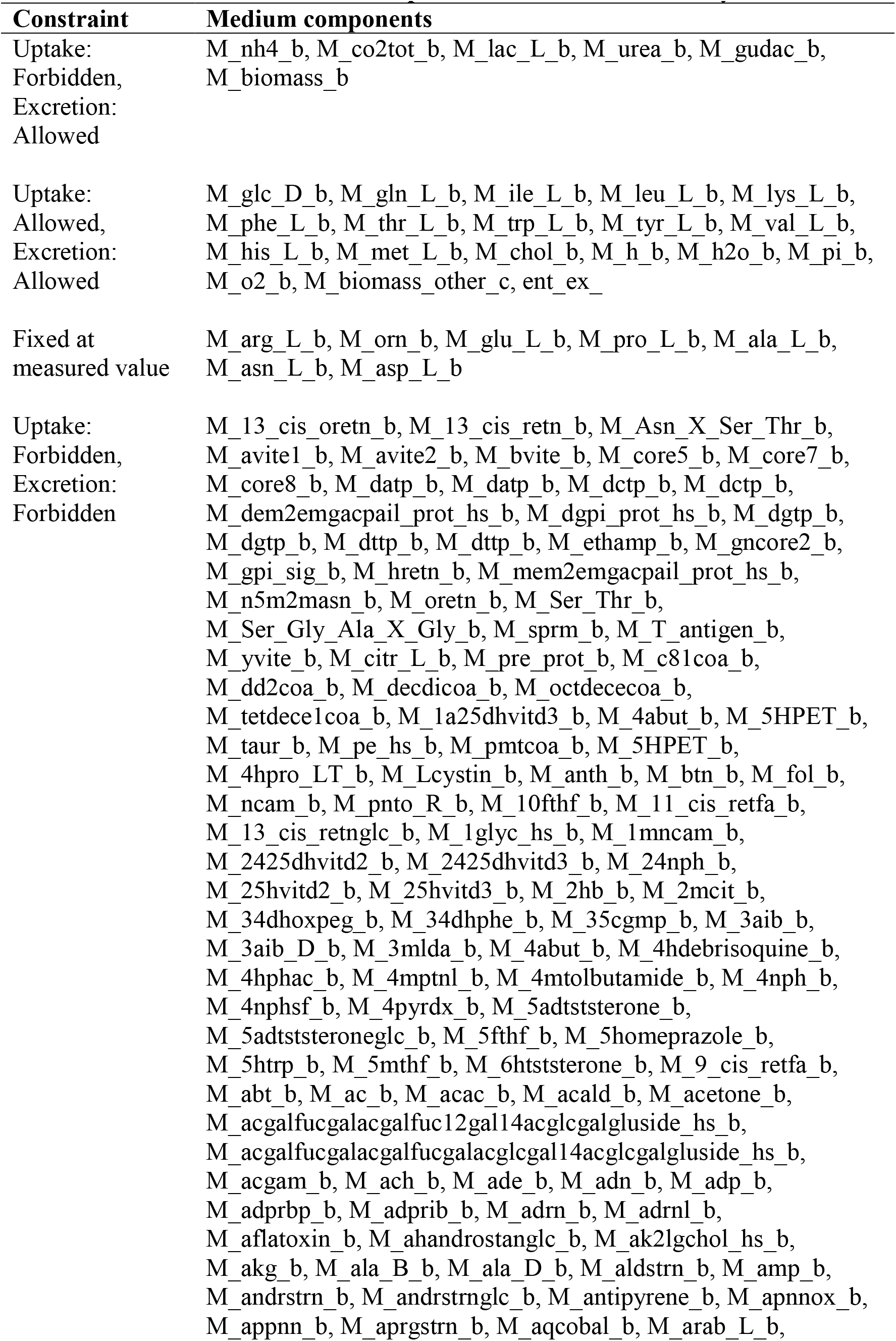
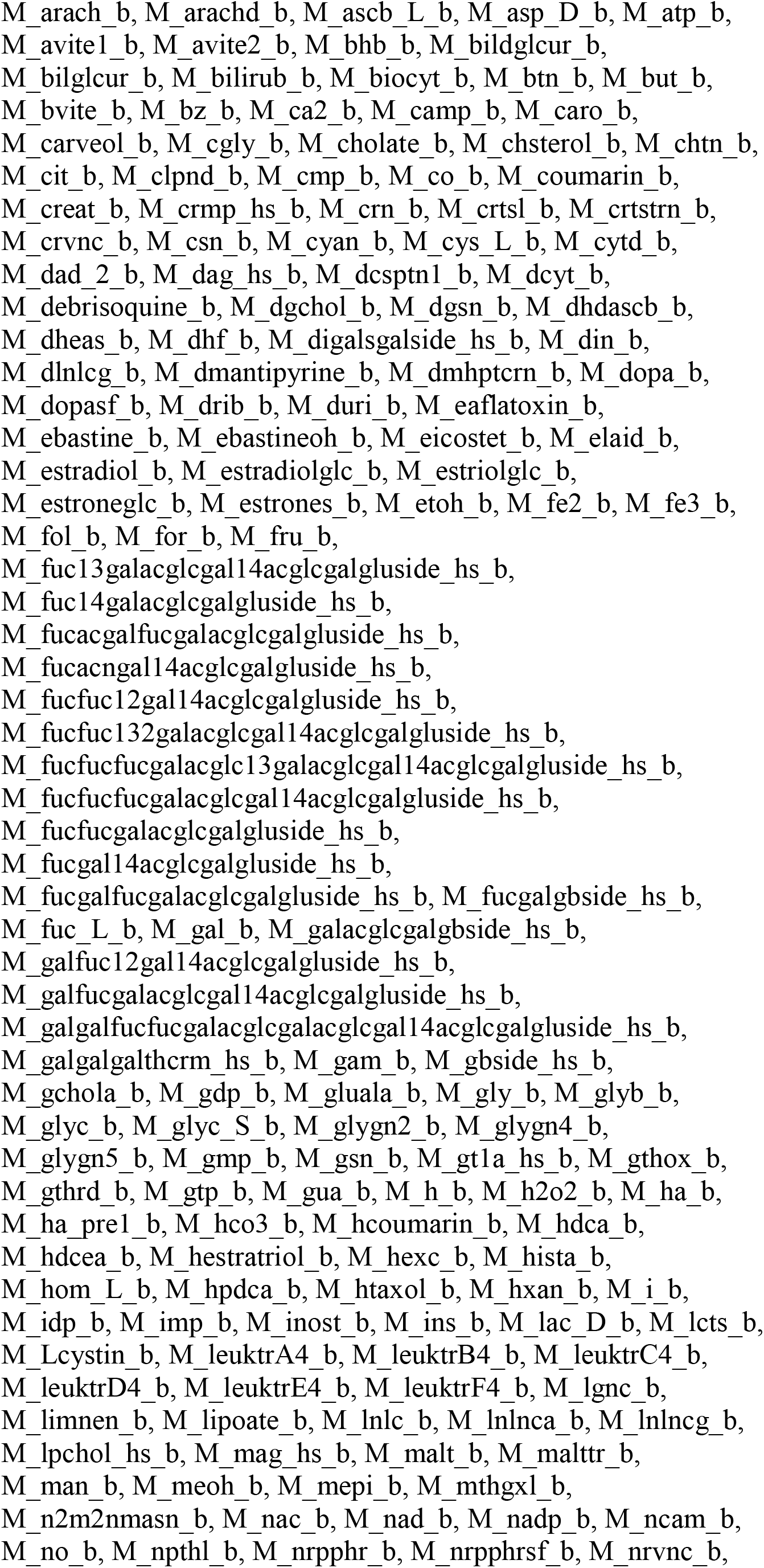
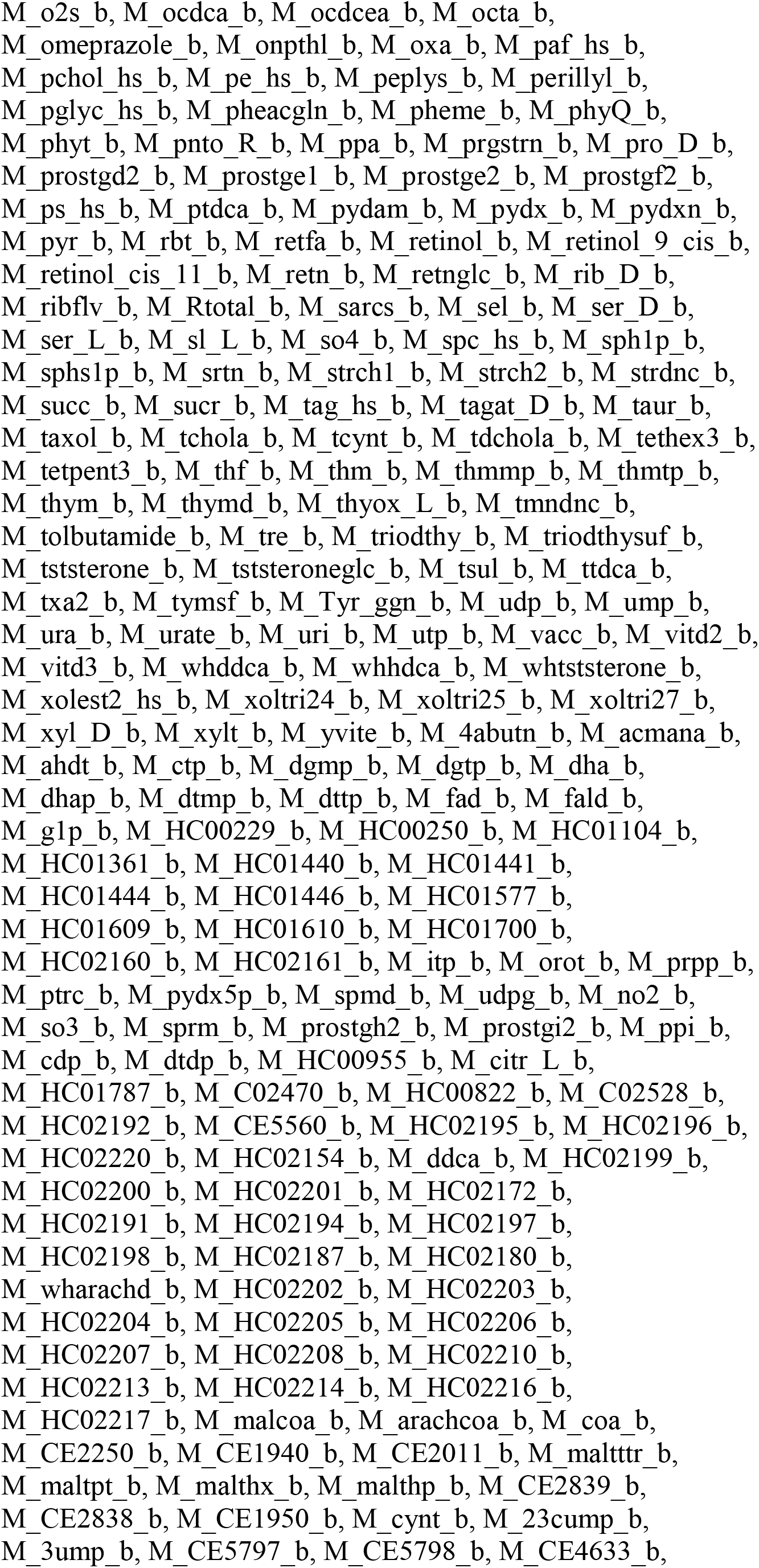
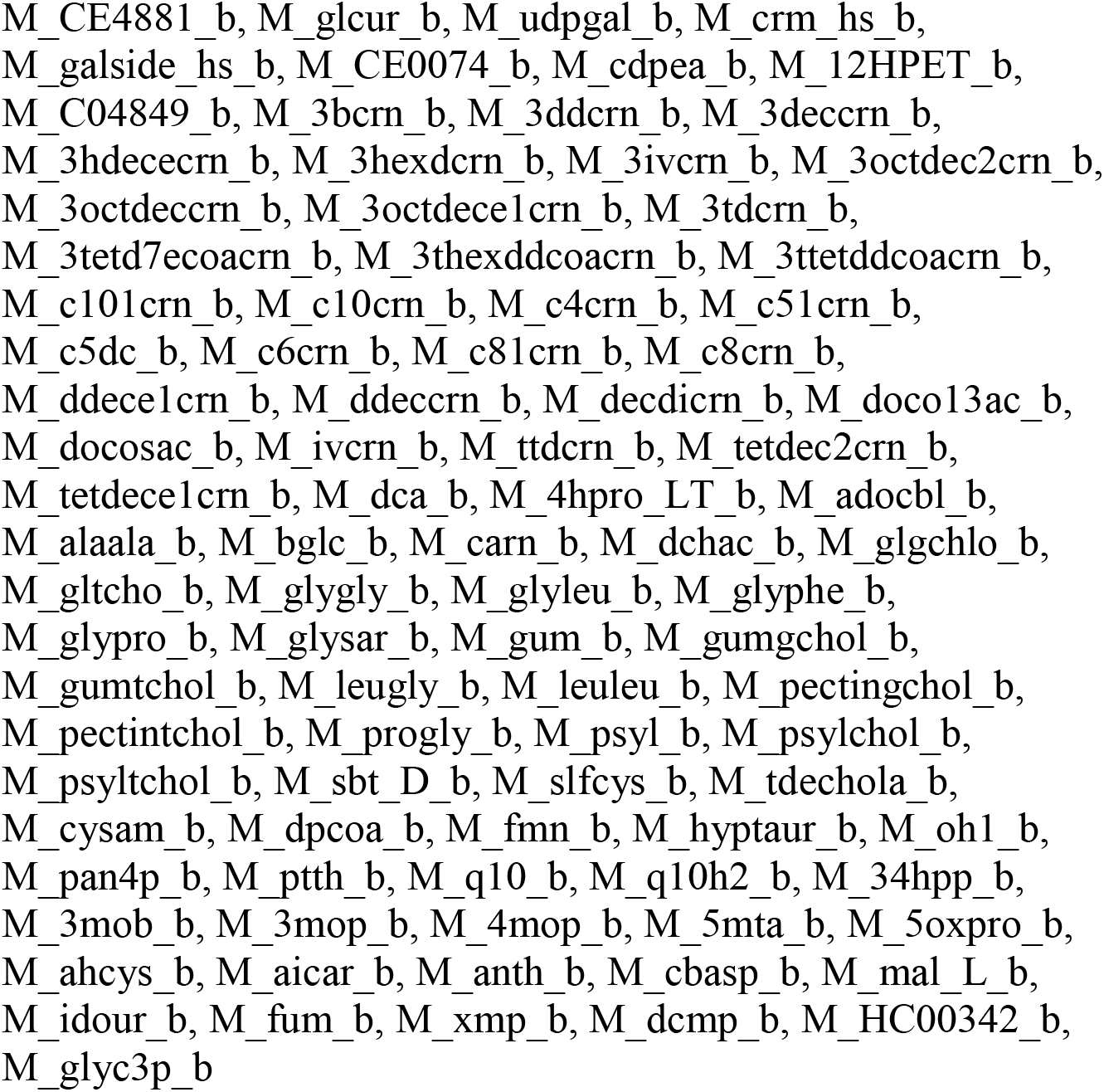
Constraints of medium components for flux balance analysis.

**Table S4.**
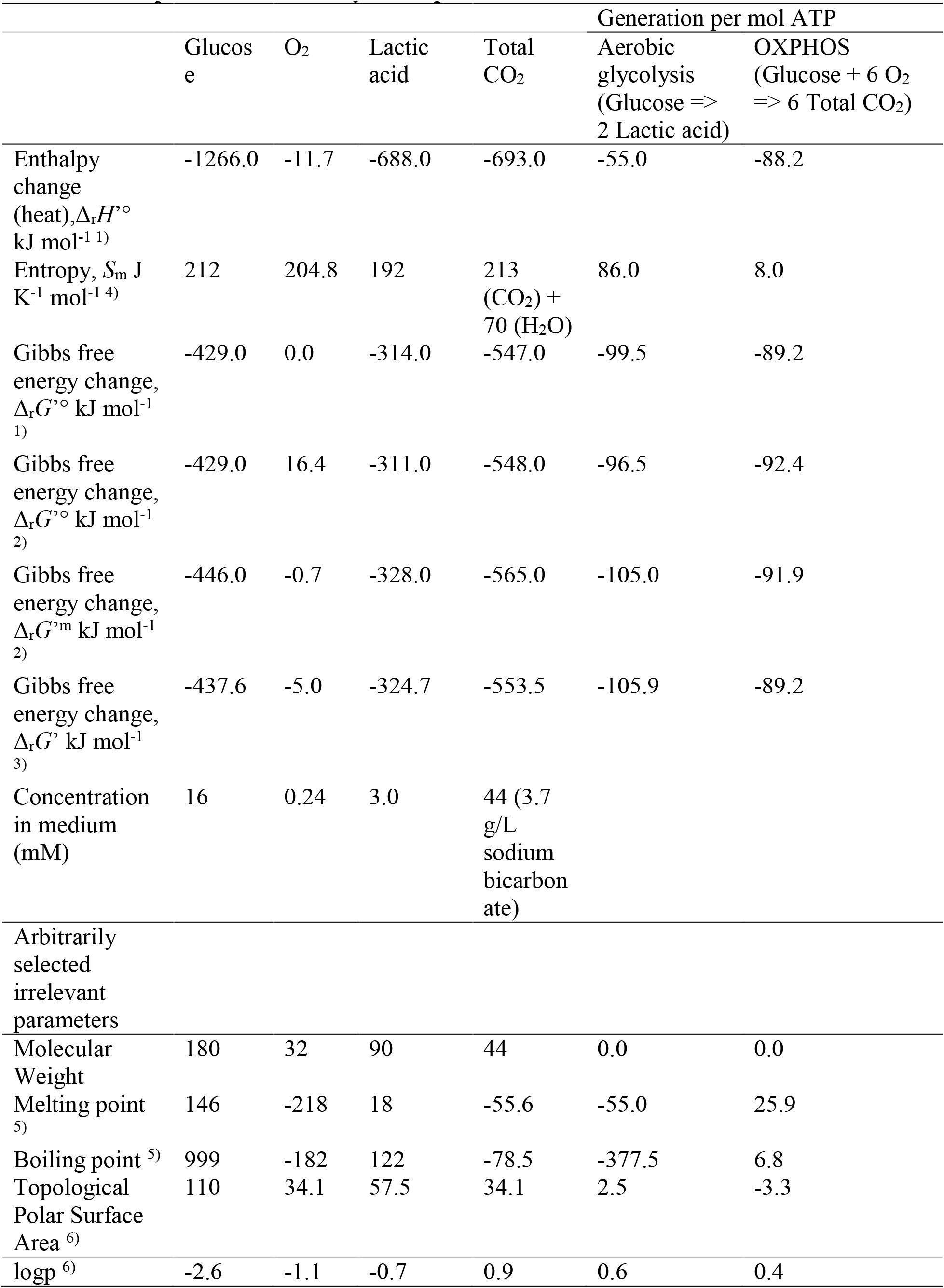
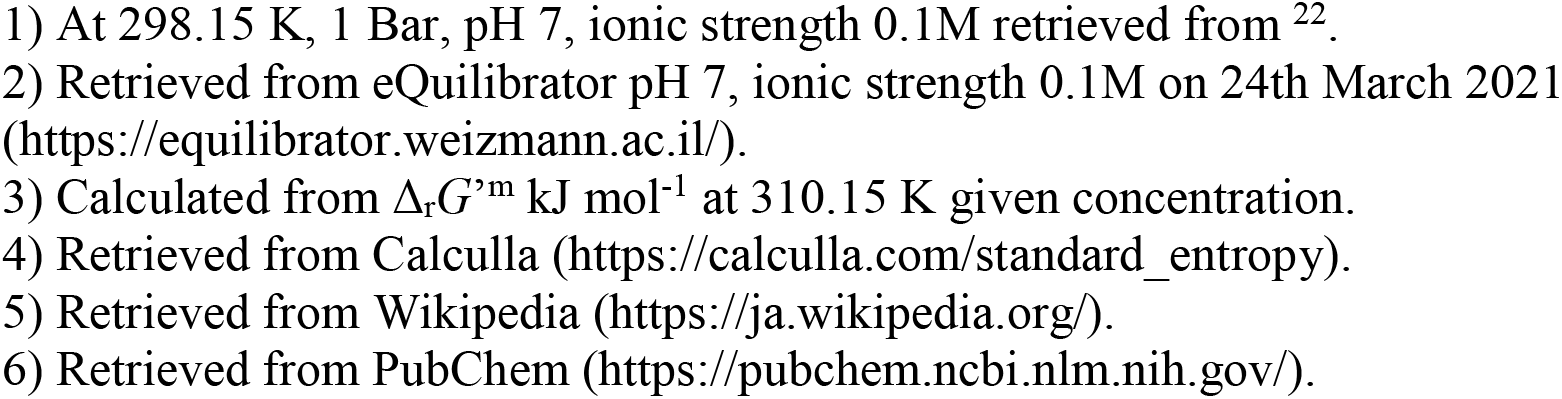
Comparison of thermodynamic parameters.

**Table S5.** Standard enthalpy of formation.

| | $\Delta_r H^\circ$ kJ mol<sup>-1</sup> 1) |
| --- | --- |
| Glucose | -1266 |
| CO<sub>2</sub> total | -693 |
| H<sub>2</sub>O | -286 |
| Lactate | -688 |
| NH<sub>4</sub><sup>+</sup>(NH<sub>3</sub>(aq)) | -133 |
| Glutamine | -808 |
| Glutamate | -982 |
| Urea | -319 |
| O<sub>2</sub> | -11.7 |
| H<sup>+</sup> | 0 |
| Alanine | -557 |
| Proline | -515 4) |
| Phosphate | -1300 |
| Arginine | -621 4) |
| Isoleucine | -638 4) |
| Leucine | -637 4) |
| Lysine | -756 5) |
| Phenylalanine | -467 4) |
| Threonine | -807 4) |
| Tryptophan | -415 4) |
| Tyrosine | -685 4) |
| Valine | -617 4) |
| Histidine | -467 4) |
| Methionine | -578 4) |
| Ornithine | -694 2) |
| Aspartate | -945 |
| Asparagine | -789.4 4) |
| Na<sup>+</sup> | -240 4) |
| Cl<sup>-</sup> | -167 4) |
| Glycine | -525 |
| Serine | -732.7 4) |
| Citrate | -1514 |
| Biomass | -5.5 kJ/g 3) |

- 1) At 298.15 K, 1 Bar, pH 7, ionic strength 0.1M retrieved from <sup>22</sup>. 2) Collected from <sup>40</sup>. 3) Collected from <sup>35</sup>. Means of yeast data were used in this study. 4) Collected from textbook (Atkins et al. Physical Chemistry for the Life Sciences) 5) Collected from <sup>41</sup>.

**Table S6.**
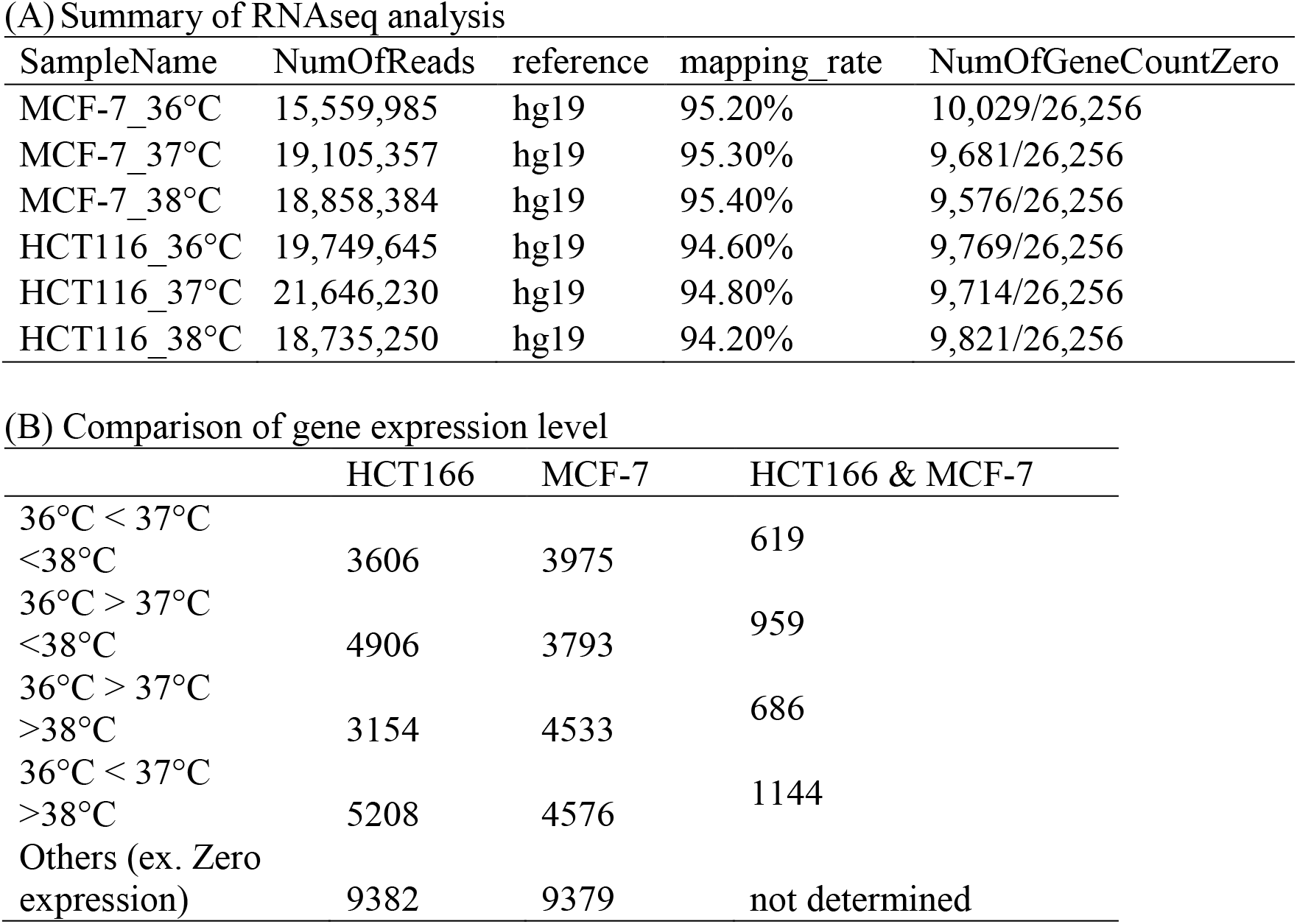
RNAseq analysis of HCT116 and MCF-7 cells cultured under different temperatures.

**Table S7.** Gene enrichment analysis using the KEGG 2019 human dataset by Enrichr.

| Term | Adjusted P-value | Genes |
| --- | --- | --- |
| Cell cycle | 1.43E-05 | PCNA;PLK1;CDC6;CDC25A;CCNA2;PTTG1;CCNE2;ORC1;TFDP2;ESPL1;ORC3;MDM2;CDK1;MCM3;ANAPC4;ANAPC5;E2F3;MCM6 |
| Proteasome | 1.43E-05 | PSMA5;PSMA6;PSMD12;PSMD6;PSMA3;PSMA2;PSMC1;POMP;PSMB1;PSME4;PSMD1 |
| Spliceosome | 2.10E-05 | TCERG1;HNRNPA3;RBM8A;DDX46;SF3B6;SRSF1;HNRNPU;WBP11;LSM3;U2SURP;HNRNPK;PRPF3;TRA2B;SRSF3;SNRPF;SRSF6;SMNDC1;RBMX |
| DNA replication | 8.40E-05 | RFC5;PCNA;RNASEH1;RFC4;LIG1;RPA1;MCM3;MCM6;DNA2 |
| Citrate cycle (TCA cycle) | 1.54E-04 | PDHA1;MDH2;IDH3G;IDH2;IDH3B;OGDHL;DLAT;SDHD |
| Oocyte meiosis | 7.20E-03 | PLK1;PPP2CA;PPP1CC;RPS6KA3;PPP3CB;PTTG1;CCNE2;ESPL1;CDK1;ANAPC4;ANAPC5;CALM1;CAMK2G |
| Mismatch repair | 2.34E-02 | RFC5;PCNA;RFC4;LIG1;RPA1 |
| Pyruvate metabolism | 4.13E-02 | ALDH3A2;LDHA;PDHA1;MDH2;DLAT;ALDH7A1 |

**Table S8.** Metabolic model for ^13^C-MFA.

| Reaction ID | Stoichiometry | Atom mapping |
| --- | --- | --- |
| r1_Glc_uptake | SubsGlc + ATP --> G6P | ABCDEF --><br>ABCDEF |
| r2_PGI_fw | G6P --> F6P | ABCDEF --><br>ABCDEF |
| r3_PGI_rv | F6P --> G6P | ABCDEF --><br>ABCDEF |
| r4_PFK | F6P + ATP --> FBP | ABCDEF --><br>ABCDEF |
| r5_FBA_fw | FBP --> DHAP + GAP | ABCDEF --> CBA +<br>DEF |
| r6_FBA_rv | DHAP + GAP --> FBP | CBA + DEF --><br>ABCDEF |
| r7_TPI_fw | DHAP --> GAP | ABC --> ABC |
| r8_TPI_rv | GAP --> DHAP | ABC --> ABC |
| r9_GAPDH_fw | GAP --> PGA + NADH + ATP | ABC --> ABC |
| r10_GAPDH_rv | PGA + NADH + ATP --> GAP | ABC --> ABC |
| r11_PEPH_fw | PGA --> PEP | ABC --> ABC |
| r12_PEPH_rv | PEP --> PGA | ABC --> ABC |
| r13_PK | PEP --> Pyr + ATP | ABC --> ABC |
| r16_PDH | Pyr --> AcCOAmit + CO2in +<br>NADH | ABC --> BC + A |
| r17_CS | AcCOAmit + Oxa --> Cit | AB + CDEF --><br>FEDBAC |
| r18_IDH_fw | Cit --> aKG + CO2in + NADH | ABCDEF --><br>ABCDE + F |
| r19_IDH_rv | aKG + CO2in + NADH --> Cit | ABCDE + F --><br>ABCDEF |
| r20_AKGDH | aKG --> Suc + CO2in + ATP +<br>NADH | ABCDE --> BCDE +<br>A |
| r21_SDH_fw | Suc-->Fum+{0.6}NADH | ABCD --> ABCD |
| r22_SDH_rv | Fum+{0.6}NADH-->Suc | ABCD --> ABCD |
| r23_FH_fw | Fum-->Mal | ABCD --> ABCD |
| r24_FH_rv | Mal-->Fum | ABCD --> ABCD |
| r25_MDH_fw | Mal --> Oxa+NADH | ABCD --> ABCD |
| r26_MDH_rv | Oxa+NADH --> Mal | ABCD --> ABCD |
| r27_PC | Pyr + CO2in + ATP --> Oxa | ABC + D --> ABCD |
| r28_MAE | Mal --> Pyr + CO2in + NADPH | ABCD --> ABC + D |
| r29_G6PDH | G6P --> m6PG + NADPH | ABCDEF --><br>ABCDEF |
| r30_6PGDH | m6PG --> Ru5P + CO2in + NADPH | ABCDEF --><br>BCDEF + A |
| r31_RPI_fw | Ru5P --> R5P | ABCDE --> ABCDE |
| r32_RPI_rv | R5P --> Ru5P | ABCDE --> ABCDE |
| r33_RPE_fw | Ru5P --> Xu5P | ABCDE --> ABCDE |
| r34_RPE_rv | Xu5P --> Ru5P | ABCDE --> ABCDE |
| r35_TKL1_fw | R5P + Xu5P --> S7P + GAP | ABCDE + FGHIJ --><br>FGABCDE + HIJ |
| r36_TKL1_rv | GAP + S7P --> Xu5P + R5P | HIJ + FGABCDE --><br>FGHIJ + ABCDE |
| r37_TAL1_fw | GAP + S7P --> F6P + E4P | ABC + DEFGHIJ --><br>DEFABC + GHIJ |
| r38_TAL1_rv | E4P + F6P --> S7P + GAP | GHIJ + DEFABC --><br>DEFGHIJ + ABC |
| r39_TKL2_fw | E4P + Xu5P --> F6P + GAP | ABCD + EFGHI --><br>EFABCD + GHI |
| r40_TKL2_rv | GAP + F6P --> Xu5P + E4P | GHI + EFABCD --><br>EFGHI + ABCD |
| r41_LDH | Pyr + NADH --> Lac | ABC --> ABC |
| r42_Lac_excretion | Lac --> LacEx | nd |
| r46_Gln_uptake | SubsGln --> Gln | ABCDE --> ABCDE |
| r47_GLC | Gln --> Glu | ABCDE --> ABCDE |
| r49_GDH_fw | Glu --> aKG | ABCDE --> ABCDE |
| r50_GDH_rv | aKG --> Glu | ABCDE --> ABCDE |
| r51_Glu_excretion | Glu --> GluEx | nd |
| r52_Pro | Glu --> Pro | ABCDE --> ABCDE |
| r53_Pro_excretion | Pro --> ProEx | nd |
| r54_Asp | Oxacyt --> Asp | ABCD --> ABCD |
| r55_Asp_excretion | Asp --> AspEx | nd |
| r56_Asn | Asp --> Asn | ABCD --> ABCD |
| r57_Asn_excretion | Asn --> AsnEx | nd |
| r58_GPT | Pyr --> Ala | ABC --> ABC |
| r59_Ala_excretion | Ala --> AlaEx | nd |
| r71_Leu_uptake | SubsLeu --> Leu | ABCDEF --><br>ABCDEF |
| r72_Ile_uptake | SubsIle --> Ile | ABCDEF --><br>ABCDEF |
| r75_Val_uptake | SubsVal --> Val | ABCDE --> ABCDE |
| r77_Arg_uptake | SubsArg --> Arg | ABCDEF --><br>ABCDEF |
| r78_Arg_degradation | Arg --> Glu | ABCDEF --><br>ABCDE |
| r79_Orn_excretion | Glu --> OrnEx | nd |
| r82_ACYL | Cit --> AcCOAcyt + Oxacyt | FEDBAC --> AB +<br>CDEF |
| r83_AcCOA_BIO | AcCOAcyt --> AcCOABiomass | nd |
| r84_G6P_BIO | G6P --> G6PBiomass | nd |
| r85_R5P_BIO | R5P --> R5PBiomass | nd |
| r86_DHAP_BIO | DHAP --> DHAPBiomass | nd |
| r87_Ser_BIO | PGA --> SerBiomass | nd |
| r89_Ala_BIO | Ala --> AlaBiomass | nd |
| r90_Asp_BIO | Asp --> AspBiomass | nd |
| r91_Asn_BIO | Asn --> AsnBiomass | nd |
| r92_Glu_BIO | Glu --> GluBiomass | nd |
| r93_Gln_BIO | Gln --> GlnBiomass | nd |
| r94_Pro_BIO | Pro --> ProBiomass | nd |
| r95_Arg_BIO | Arg --> ArgBiomass | nd |
| r96_Leu_BIO | Leu --> LeuBiomass | nd |
| r97_Ile_BIO | Ile --> IleBiomass | nd |
| r98_Val_BIO | Val --> ValBiomass | nd |
| r100_CO2_in | CO2in --> CO2Ex | A --> A |
| r101_CO2_ex | SubsCO2 --> CO2in | A --> A |
| r104_MAL_transport_fw | Mal --> Malcyt | ABCD --> ABCD |
| r105_MAL_transport_rv | Malcyt --> Mal | ABCD --> ABCD |
| r106_MDH_cyt_fw | Malcyt --> Oxacyt + NADH | ABCD --> ABCD |
| r107_MDH_cyt | Oxacyt + NADH --> Malcyt | ABCD --> ABCD |
| r108_Net_ATP_production | ATP --> ATPex | nd |
| r109_OXPHOS | NADH --> 2.5 ATP | nd |
| r110_Net_NADPH_production | NADPH --> NADPHex | nd |

